# Serum Amyloid P Secreted by Bone Marrow Adipocytes Drives Skeletal Amyloidosis

**DOI:** 10.1101/2024.08.15.608092

**Authors:** Surendra Kumar, Kangping Song, Jiekang Wang, Meghraj Singh Baghel, Philip Wong, Xu Cao, Mei Wan

## Abstract

The accumulation of amyloid fibrils has been identified in tissues outside the brain, yet little is understood about the formation of extracerebral amyloidosis and its impact on the aging process of these organs. Here, we demonstrate that both transgenic mice modeling Alzheimer’s disease (AD) and naturally aging mice exhibit accumulated senescent bone marrow adipocytes (BMAds), accompanied by amyloid deposits surrounding the BMAds. Senescent BMAds acquire a secretory phenotype, resulting in a marked increase in the secretion of serum amyloid P component (SAP), also known as pentraxin 2 (PTX2). SAP/PTX2 colocalizes with amyloid deposits around senescent BMAds *in vivo* and is sufficient to promote the formation of insoluble amyloid deposits from soluble Aβ peptides in *in vitro* and *ex vivo* 3D BMAd-based culture experiments. Additionally, Combined treatment with SAP/PTX2 and Aβ peptides promotes osteoclastogenesis but inhibits osteoblastogenesis of the precursor cells. Transplantation of senescent BMAds into the bone marrow cavity of healthy young mice is sufficient to induce bone loss. Finally, pharmacological depletion of SAP/PTX2 from aged mice abolishes bone marrow amyloid deposition and effectively rescues the low bone mass phenotype. Thus, senescent BMAds, through the secretion of SAP/PTX2, contribute to the age-associated development of skeletal amyloidosis and resultant bone deficits.

## INTRODUCTION

Alzheimer’s disease (AD), the most prevalent form of dementia, is characterized by the progressive accumulation of insoluble amyloid-β (Aβ) plaques in the brain parenchyma and vasculature, along with neurofibrillary tangles of hyperphosphorylated tau, resulting in brain degeneration and cognitive decline ^1–3^. Although the primary symptoms of amyloid Aβ deposits predominantly affect the central nervous system, increasing evidence has demonstrated that insoluble amyloid fibril can form and deposit in other organ systems. Indeed, amyloid fibril aggregation have been captured in several non-brain organs, such as the heart, kidneys, liver, lungs, peripheral nerves, and gastrointestinal tract ^4–6^. In fact, systemic amyloidosis typically describes a set of rare conditions characterized by the buildup of amyloid fibrils in various organs and tissues throughout the body ^7^. The common etiology of various amyloid diseases remains unclear at present. While gene mutations have been identified in rare hereditary amyloidosis, disrupted protein homeostasis associated with aging is likely a primary pathogenic factor for amyloid deposits in general ^8^.

Amyloidosis damage tissue and organs through deposition of misfolded protein form “amyloid fibrils” in their extracellular space. Amyloid fibrils are structured aggregates of polypeptides with a cross-β structure ^9–11^. To date, 42 different amyloid proteins have been identified, each associated with specific clinical forms of amyloidosis ^12,13^. Most types of amyloidosis are age-associated. Over ten amyloid fibrils, such as tau, Aβ, and wild-type transthyretin in humans and apolipoprotein A-II in mice, have been linked to disease development ^14^. It is known that amyloid fibrils are formed from Aβ peptide, which is produced at cholesterol-rich regions of neuronal membranes and secreted into the extracellular space ^10,15^. Aβ peptides have tendency to aggregate in different shapes such as fibrillar structure “amyloid fibrils” ^16,17^ and /or non-fibrillar accumulates which also known as “ Aβ oligomers”^18,19^. Amyloid fibril formation is driven by several factors, including increased protein concentration, misfolding triggers, and proteolytic modification into amyloidogenic fragments. These fibrils interact with serum amyloid P component/pentraxin 2 (SAP/PTX2), apolipoprotein E, and glycosaminoglycans, which are crucial for amyloid deposit assembly and maintenance in tissues.^20–24^. However, despite extensive research on the role of intracerebral amyloid deposition in neurodegeneration within central nervous system, the formation of extracerebral amyloidosis, its biochemical and pathological features, and its impact on the aging process of other organs have been little studied so far.

Skeletal health holds a unique connection to AD. Patients with osteoporosis often have a higher incidence of AD, and bone loss is frequently detected early in the progression of AD, preceding significant cognitive decline ^25–27^. Notably, low bone mineral density (BMD) can predict risk of AD years before dementia diagnosis ^26,28,29^. Similar to this clinical comorbidity, reduced BMD traits are exhibited in various transgenic mouse models designed to mimic humanized Alzheimer’s disease neuropathology ^30,31^. These findings suggest that age-associated skeletal pathologies likely involve mechanisms directly associated with AD pathology. We postulate that amyloid fibrils may also form and deposit in bone tissue similarly to those in brain tissue. While amyloid depositions have been reported in several organs of aged patients such as knee joints, intervertebral discs, and ligamentum flavum ^32–34^, there has been no investigation into whether amyloidosis also occurs in bone/bone marrow with age or under conditions of AD, and how it may contribute to any pathologic consequences in the skeleton. Here, we show that amyloid fibrils deposit in bone marrow of both AD mice and aged mice, both of which exhibit a phenotype of decreased bone mass and increased marrow adiposity. Using various *in vivo* and *in vitro* approaches, we identify senescent bone marrow adipocytes (BMAds) as a major culprit for the formation of bone amyloid deposition. Senescent BMAds, driven by CEBPα-activated *p19Arf*, secrete SAP/PTX2, thereby inducing amyloid deposition and leading to altered osteoblast and osteoclast activities for bone impairment.

## RESULTS

### Amyloid deposition is accumulated in bone/bone marrow of AD mice and nature aging mice

To unravel the mechanisms contributing to bone deficits in AD, we first examined the skeletal phenotype of *APPswe/PS1dE9* (*APP-PS1*) transgenic mice, a well-established AD mouse model ^35,36^. At nine months of age, *APP-PS1* mice (named “AD” hereafter) displayed a low bone mass phenotype (Figure 1A), characterized by a significant decrease in bone volume per tissue volume (BV/TV%) (Figure 1B), trabecular thickness (Tb.Th) (Figure 1C), and trabecular number Tb.N) (Figure 1D), along with an increase in trabecular separation (Tb.Sp) (Figure 1E) compared to age-matched wild-type littermates (WT). Cortical bone analysis revealed an unchanged cortical area (Ct.Ar) (Figure 1G) but a reduced cortical thickness (Ct.Th) (Figure 1H), reflecting age-associated cortical bone changes. A similar bone loss phenotype was observed in 5xFAD (Figure S1), another widely used AD transgenic mouse model ^37,38^. Additionally, there was greater bone marrow adipose tissue mass (Figure 1I-J), with markedly higher number and cell size of Perilipin^+^ adipocytes (Figure 1K-M) in AD mice relative to WT mice. Since aging-related osteoporosis typically features low bone mass, microarchitectural alterations, and bone marrow adipose tissue expansion ^39–41^, our findings suggest that AD mice at a younger age recapitulate an aged skeletal phenotype.

**Figure 1.**
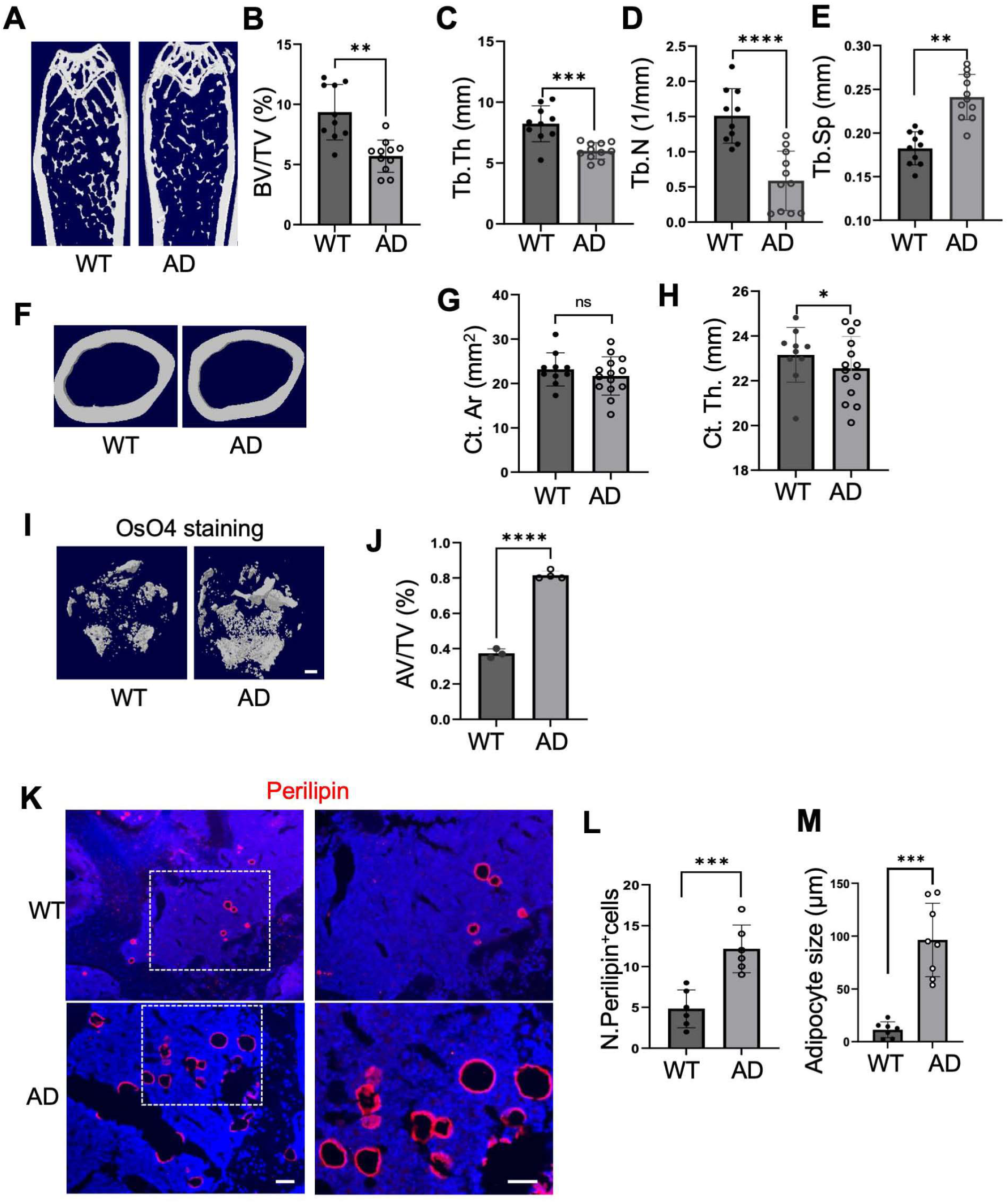
*APP-PS1* mice recapitulate an aged bone phenotype. Femoral and tibial bones were harvested from 9-month-old male and female mice carrying the *APP_swe_/PS1_ΔE9_ (APP-PS1)* transgene, referred to as “AD” hereafter, for bone phenotypic assessments. (**A-H**) Representative μCT images of the trabecular area of the distal femur are shown in (**A**) with quantitative analyses of trabecular bone volume (BV/TV) (**B**), trabecular thickness (Tb. Th) (**C**), trabecular number (Tb. N) (**D**), and trabecular separation (Tb. Sp) (**E**) n=10-11. Representative μCT images of the cortical area of the femur are shown in (**F)** with quantitative analyses of cortical area (Ct. Ar) **(G)** and cortical thickness (Ct. Th) **(H)**. n=10-11. **(I, J)** Representative μCT images of OsO4-stained decalcified distal femurs are shown in (I) with quantitative analysis of adipose tissue volume per total volume (% AV/TV) in (J). Scale bar = 500 pm. n=4 mice. **(K-M)** Immunofluorescence staining of tibia bone sections was performed using antibody against Perilipin. Representative images of Perilipin^+^ cells from secondary spongiosa area of tibia are shown in (**K**). Analyses of the cell number per mm^2^ tissue area (N. Perilipin^+^ cells) and the cell size of the adipocytes (pm) are shown in (**L**) and (**M**), respectively. Scale bar = 200 pm. n=6-8. Data are represented as mean ±s.e.m. *p< 0.05, **p<0.01, ***p<0.001, as determined by unpaired two-tailed Studenf s t-tests for 2 groups.

Age-related amyloidosis has been identified in several tissues outside of the brain. We examined whether Aβ fibrils are also present in the bone and bone marrow of AD mice and nature aging mice. Quantification by ELISA revealed increased Aβ1-40 and Aβ1-42 levels in the bone marrow of *APP-PS1* mice compared to the WT mice (Figure 2A-B). Intriguingly, both Aβ1-40 and Aβ1-42 levels were also markedly elevated in the bone marrow of 22-month-old (Old) compared to 4-month-old mice (Young) (Figure 2C-D). Dot blot filter retention assays, a technique that separates insoluble Aβ aggregates from soluble Aβ by retention on a filter membrane, confirmed a significant increase in Aβ oligomers in the bone marrow of AD mice and old mice compared to the WT littermates and young mice, respectively (Figure 2E). We then attempted to visualize the presence of Aβ deposits in bone tissue of AD mice using thioflavin-S (ThioS) staining and Aβ amyloid immunostaining. While the WT mice had no obvious Aβ deposits in bone and bone marrow, extensive small ThioS^+^ fibrillar amyloid deposits were observed in AD mice (Figure 2F-G). Immunostaining of the bone tissue sections with H31L21, an antibody recognizing both human and mouse APP, showed fewer but larger Aβ plaques than ThioS staining in bone/bone marrow of AD mice (Figure 2H-I). We also evaluated if fibrillar amyloid deposits occur in bone/bone marrow of nature aging mice. Like the finding in AD mice, extensive but diffuse ThioS^+^ fibrillar aggregations were found in the bone and bone marrow of the mice at 22 months of age (Old) with no Aβ amyloid deposits in the same region of 4-month-old mice (Young) (Figure 2J-K). In the Old mice, increased H31L21^+^ deposits were extensive but generally smaller than those in AD mice (Figure 2L-M). A notable observation is that a significant portion of diffuse H31L21^+^ deposits formed a large, ring-like structure around BMAds (Figure 2L, lower panel), indicating that BMAds may possess the ability to stabilize Aβ fibrils in their vicinity through certain mechanisms.

**Figure 2.**
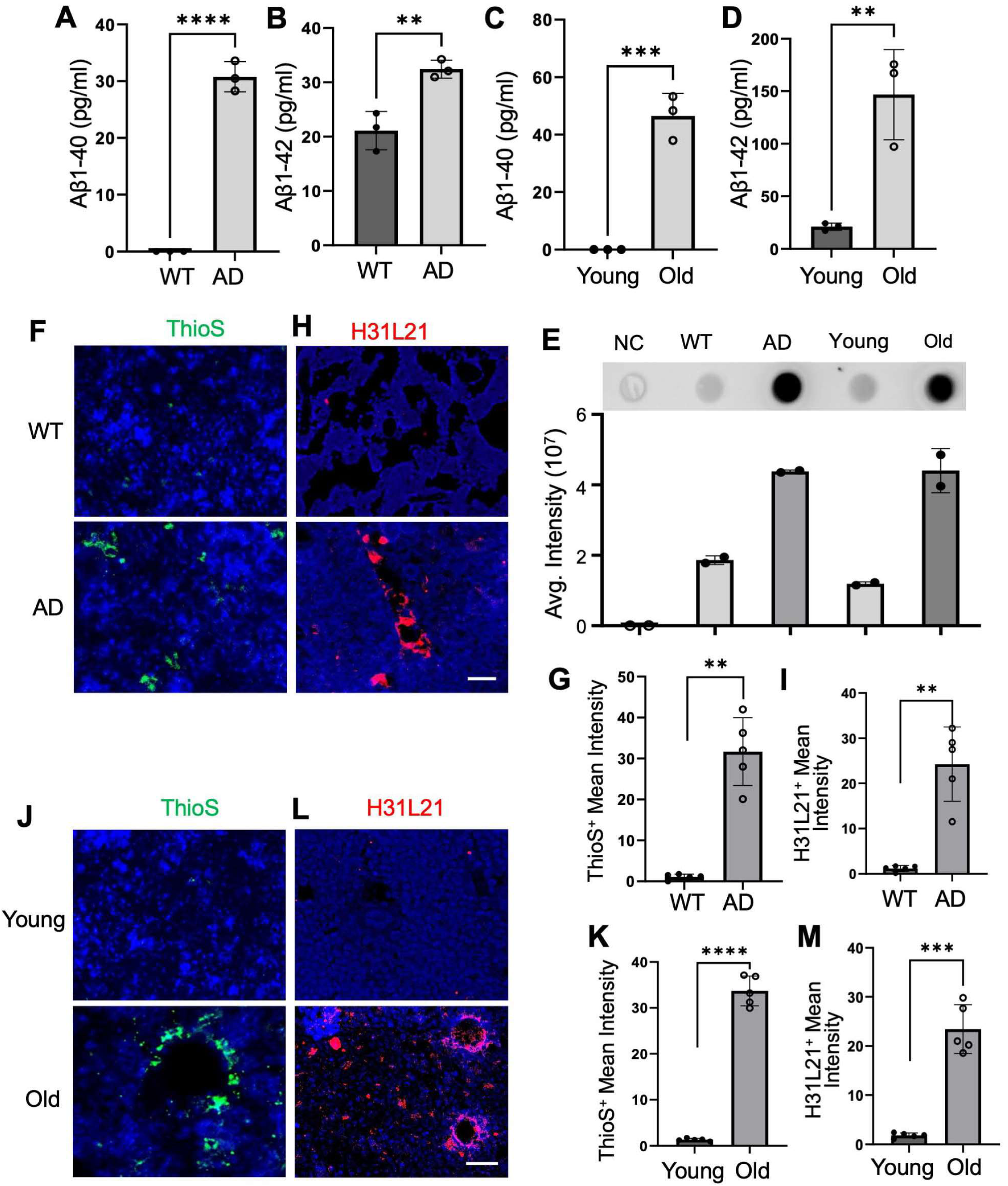
AD mice and nature aging mice develop amyloid deposition around BMAds. (**A-D**) Concentrations of Aβ peptides (Aβ1-40 and Aβ1-42) in bone marrow were measured by ELISA in 9-month-old male *APP-PS1* mice (AD) compared to wild-type littermates (WT) (**A** and **B**), and in 24-month-old male C57BL/6 mice (old) compared to 4-month-old male C57BL/6 mice (Young) (**C** and **D**). n= 3. (**E**) Bone marrow was collected from different mouse strains as indicated. Amyloid aggregates were detected by filter retention assays. A representative image is shown in the upper panel with quantified values shown in lower panel. (**F-I**) Tibial bones were harvested from 9-month-old *APP- PS1* mice (AD) and their wild-type littermates (WT). Aβ deposits were detected by staining frozen bone sections with ThioflavinS (ThioS) (**F**) and by immunofluorescence staining of paraffin bone sections with antibody against H31L21 (**H**). Quantification of ThioS^+^ mean intensity and H31L21^+^ mean intensity was shown in (**G**) and (**I**), respectively. Scale bar = 200 pm. n= 5. (**J-M**) Tibial bones were harvested from 24-month-old C57BL/6 mice (Old) compared to 4-month- old C57BL/6 mice (Young). Aβ deposits were detected by staining of frozen bone sections with ThioS (**J**) and by immunofluorescence staining of paraffin bone sections with antibody against H31L21 (**L**). Quantification of ThioS^+^ mean intensity and H31L21^+^ mean intensity was shown in (**K**) and (**M**), respectively. Scale bar = 200 pm. n= 5. Data are represented as mean ±s.e.m. **p<0.01, ***p<0.001, ****p<0.0001, as determined by unpaired two-tailed Student’s t-tests for 2 groups.

### BMAds undergo cellular senescence mediated by CEBPα-activated *p19Arf*

We recently found that BMAds undergo rapid cellular senescence and acquire a SASP in mice following glucocorticoid treatment ^42^. To investigate whether BMAds also exhibit senescence phenotype in AD mice and nature aging mice, we first assessed SA-βGal expression. In 9-month-old WT mice, SA-βGal^+^ cells were sparsely distributed in bone/bone marrow, while same age of AD mice exhibited higher abundance of SA-βGal^+^ cells. When bone tissue sections were co­stained with SA-βGal and Oil Red O, it was evident that AD mice exhibited notably higher levels of Oil Red O-positive BMAds and a greater percentage of SA-βGal^+^ BMAds compared to their WT littermates (Figure S2A-C), indicating a senescence-like change in BMAds under AD conditions. We then examined the RNA expressions of *p16Ink4a* and *p21Cip*, two cellular senescence hallmarks, in bone tissue of AD mice by RNAscope analysis. Both *p16Ink4a* and *p21Cip* expression were almost absent in bone/bone marrow of WT littermate control mice, whereas *p16Ink4a*^+^ (Figure S2D-E) and *p21Cip*^+^signals (Figure S2F-G) were observed in the vicinity of greater numbers of BMAd-like cells in AD mice compared to WT littermates. As *p19Arf* is the upstream activator of the p53-p21 pathway, we assessed the mRNA expression of *p16Ink4a*, *p19Arf*, and *p21Cip* by qRT-PCR. As expected, the expressions of all three senescence genes were significantly elevated in bone marrow of AD mice compared to their WT littermates (Figure S2H).

Further RNAscope analysis of bone tissue sections revealed that the *p19Arf*^+^ signal was almost exclusively localized around large, ring-like BMAds in both AD mice (Figure 3A-B) and Old mice (Figure 3D-E). In contrast, no *p19Arf*^+^ signal was detected in WT littermates and young mice, respectively. Additionally, we observed increased *H2AX*^+^ signals, indicative of reactive oxygen species (ROS) generation, DNA damage and repair ^43,44^. The increased *H2AX*^+^ signals overlapped with *p19Arf*^+^ signal around BMAds in both AD mice (Figure 3A, C) and old mice (Figure 3D, F). Double-immunofluorescence staining of bone tissue sections confirmed a significant increase in the protein expression of the *p19Arf* in Perilipin^+^ BMAds in AD mice (Figure 3G-H). Immunofluorescence using the DNA damage marker γH2AX, a robust marker of senescent cells ^45–47^, also revealed a substantial increase in γH2AX+ BMAds in both AD mice (Figure 3I-J) and Old mice (Figure 3K-L) compared with their respective controls.

**Figure 3.**
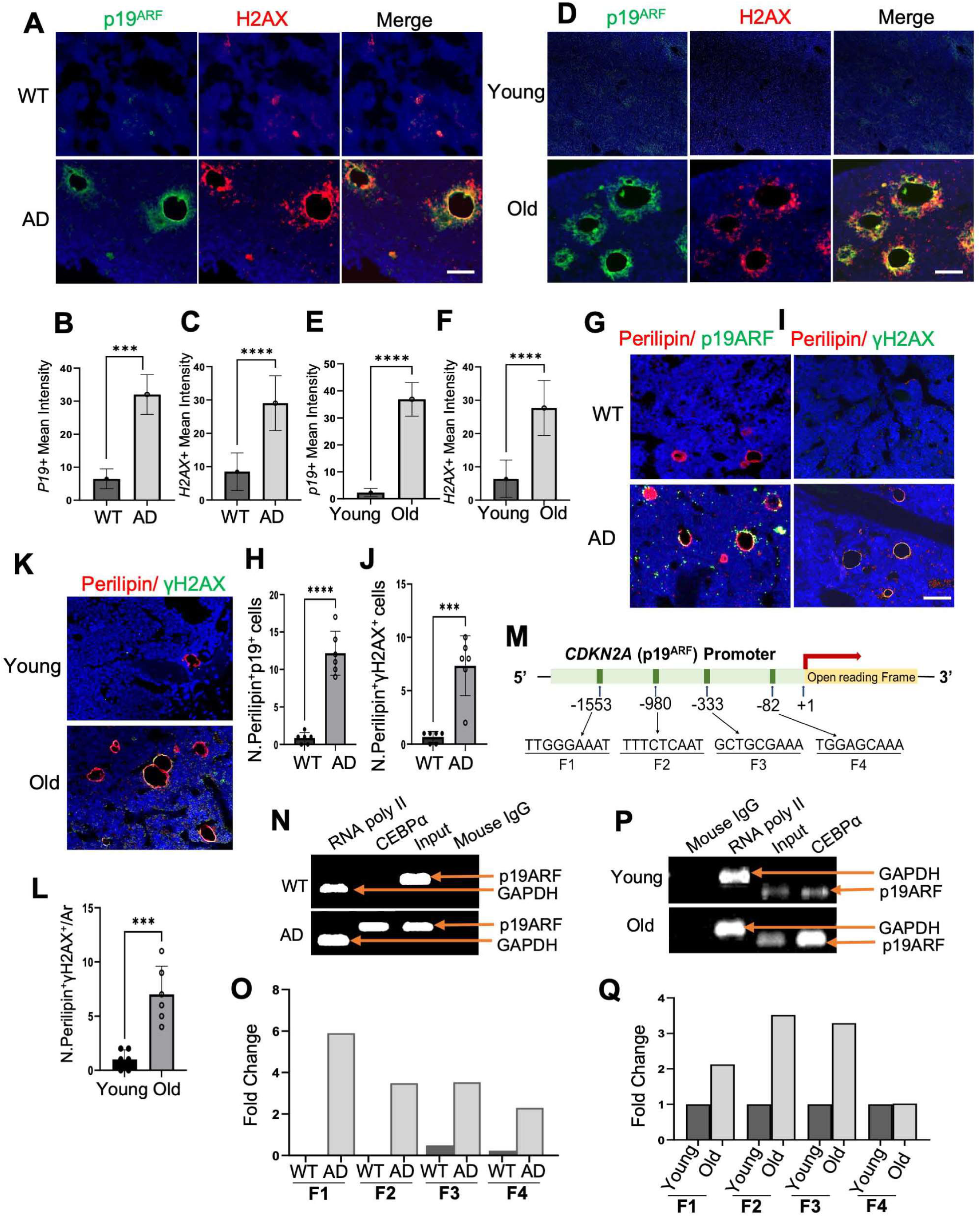
BMAds undergo cellular senescence mediated by CEBPα-activated *p19Arf*. (**A-C**) Tibial bones were harvested from 9-month-old *APP-PS1* mice (AD) and their wild-type littermates (WT). Bone tissue sections were subjected to RNAscope assays using specific probes to *p19Arf* and *H2AX*. Representative images are shown in figure (**A**). The quantification of p19Arf^+^ mean intensity and H2AX^+^ mean intensity was shown in (**B**) and (**C**), respectively. n= 6. (**D-F**) Tibial bones were harvested from 24-month-old C57BL/6 mice (Old) compared to 4-month- old C57BL/6 mice (Young). Bone tissue sections were subjected to RNAscope assays using specific probes to *p19Arf* and *H2AX*. Representative images are shown in figure (**D**). Quantification of *p19Arf ^+^* mean intensity and *H2AX* ^+^ mean intensity is shown in (**E**) and (**F**), respectively. Scale bar = 100 pm. n= 6. (**G-J**) Double-immunofluorescence staining of tibial bone sections from *APP-PS1* mice (AD), and wild-type littermates (WT) was performed using antibodies against Perilipin and p19Arf or γH2AX. Representative images of Perilipin^+^p19Arf^+^ cells from secondary spongiosa area of tibia are shown in (**G**), and analysis of cell number per mm^2^ tissue area (N. Perilipin^+^p19Arf^+^ cells) are shown in (**H**). Representative images of Perilipin+γH2AX ^+^ cells from secondary spongiosa area of tibia are shown in (**I**), and analysis of cell number per mm^2^ tissue area (N. Perilipin^+^γH2AX ^+^ cells) are shown in (**J**). Scale bar = 200 pm. n=6. (**K-L**) Double-immunofluorescence staining of tibia bone sections from 24-month-old mice (Old) and 4-month-old mice (Young) was performed using antibody against Perilipin and γH2AX. Representative images of Perilipin^+^p19Arf^+^ cells from secondary spongiosa area of tibia are shown in (**K**), and analysis of cell number per mm2 tissue area (N. Perilipin^+^ γH2AX ^+^ cells) are shown in (**L**). Scale bar= 200 pM. n=6. (**M-Q**) Chromatin immunoprecipitation (ChIP)-qPCR assays. Schematic diagram of the *p19Arf* promoter with four CEBPα-binding sites (M). Chromatin immunoprecipitation (ChIP) assay on the promoter of *p19Arf* in isolated BMAds from *APP-PS1* mice (AD) and wild-type littermates (WT) (**N-O**) and from 24-month-old mice (Old) and 4-month-old mice (Young) (**P-Q).** CEBPα antibody or IgG antibody (negative control) were used for the ChIP assay. ChIP and input DNA were measured using RT-qPCR. RNA poly II was included as a positive control. Representative agarose gel pictures are shown in (**N**) and **(P**). Quantitively fold enrichment ratios for DNA immunoprecipitated by CEBPα at different binding sites are shown in (O) and (Q). Data are represented as mean ±s.e.m. ***p<0.001, ****p<0.0001, as determined by unpaired two-tailed Student’s t-tests for 2 groups.

We then investigated the mechanisms underlying *p19Arf* activation in BMAds during aging or under AD conditions. Although they have largely overlapping DNA sequences, the *INK4a-ARF* locus produces two separate proteins: p16Ink4a and p19Arf. Analysis of the *INK4a-ARF* locus identified four putative CEBPα and two putative PPARγ binding sites at the promoter regions of both *p16In4a* (Figure S3A) and *p19Arf* (Figure 3M). These putative binding sites are distinct between *p16Ink4a* and*p19Arf* promoters. We previously found that PPARγ, a master regulator of adipogenesis, mediates glucocorticoid-induce BMAd senescence characterized by *p16Ink4a* activation ^42^, and PPARγ can bind to the *p16Ink4a* promoter and induce its transcription ^48^. To assess the status of CEBPα, a pleiotropic transcriptional activator known to promote adipogenesis, along the promoters of both *p16Ink4a* and *p19Arf* in bone marrow cells, we conducted ChIP assays. No specific CEBPα bindings to the four CEBPα binding sites were detected in the promoter of *p16Ink4a* in bone marrow cells of either WT or AD mice (Figure S3B). Intriguingly, enriched CEBPα was detected in all four regions in the promoter of *p19Arf* in bone marrow cells of AD mice but not in WT mice (Figure 3N-O). Similarly, CEBPα was significantly more enriched at the three of four putative binding sites in the *p19Arf*promoter in bone marrow cells of old mice relative to young mice (Figure 3P-Q). Taken together, the results suggest that CEBPα-activated *p19Arf* gene is a key mediator of BMAd senescence during aging and under AD conditions.

### Senescent BMAds is sufficient to induce low bone mass phenotype by producing SASP factors

An important feature of senescent cells is SASP, through which these cells can exert detrimental effects on their surrounding microenvironment ^49–52^. To investigate whether senescent BMAds acquire a SASP, we conducted proteomics assays. BMAds isolated from AD mice and age-, sex-matched WT littermates were cultured using the approach illustrated in Figure 4A. Conditioned media (CM) from BMAds were collected and subjected to antibody array screening analyses to identify secreted factors. Seventeen out of 111 adipokines/cytokines/growth factors were found to be increased in the CM of BMAds isolated from AD mice compared to those from WT mice (blue and red squares in Figure 4B-C). Notably, SAP/PTX2 exhibited the highest increase among the identified factors (red square in Figure 4B-C), indicating its role as one of the most prominently released SASP factors from senescent BMAds.

**Figure 4.**
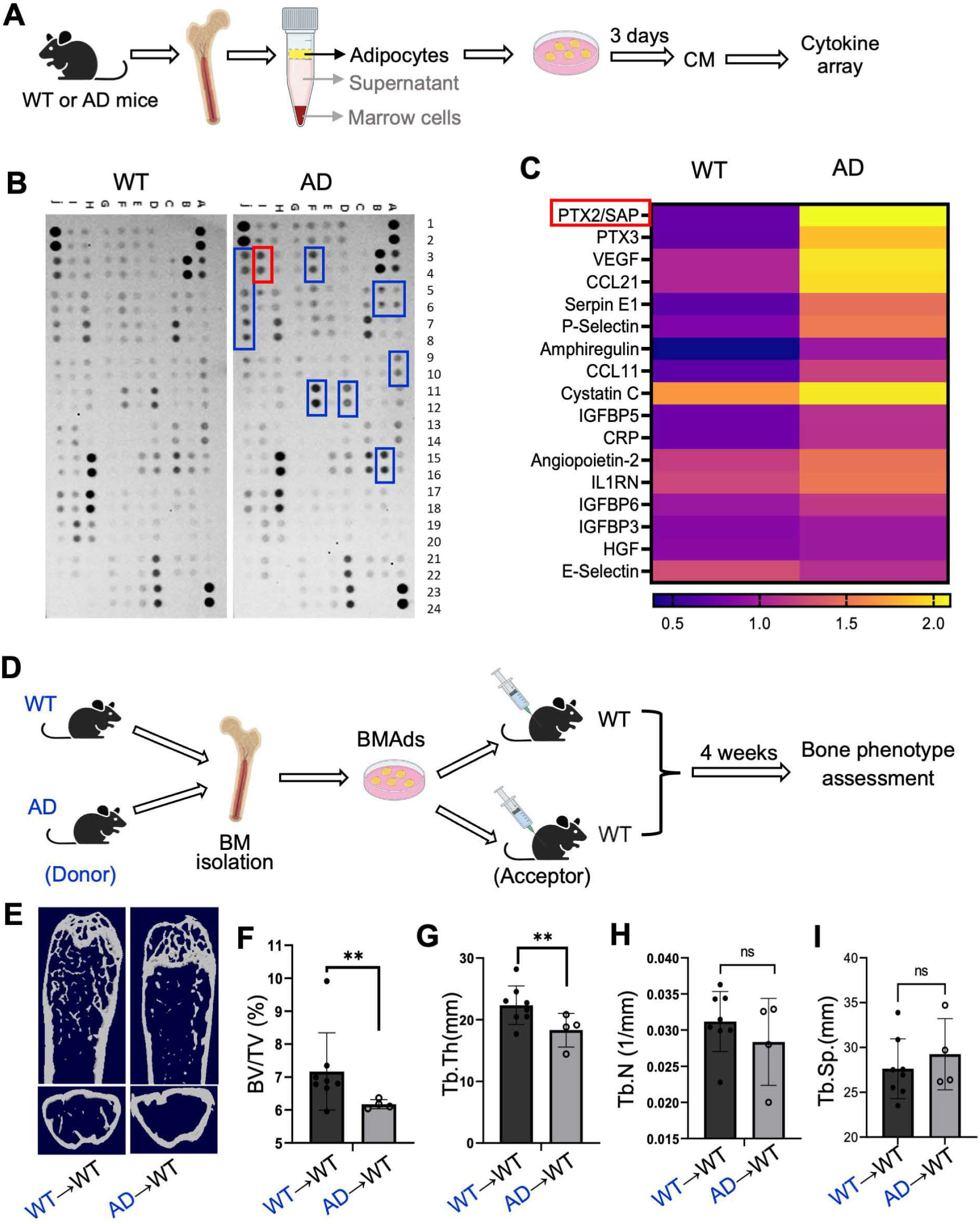
Senescent BMAds are sufficient to induce low bone mass phenotype by producing SASP factors. (**A**) A schematic diagram illustrating the procedure of the conditioned medium (CM)-based cytokine array assays. BMAds were isolated from 9-month-old *APP-PS1* mice (AD) and wild-type littermates (WT), cultured in alpha-minimal essential medium (aMEM) with 1% PS for 3 days. **(B- C**) Secreted proteins in the CMs were characterized using cytokines/adipokines arrays (**B**). HeatmAβ of differentially expressed cytokines in the CMs is shown in (**C**). (**D-I**) Schematic diagram showing the experimental design for transplantation (**D**). Representative μCT images of femoral bones are shown in (**E**) with quantitative analyses of trabecular bone volume/tissue volume (BV/TV) (**F**), trabecular thickness (Tb. Th) (**G**), trabecular number (Tb. N) (**H**), and trabecular separation (Tb. Sp) (**I**). n = 4-8. Data are represented as mean ± SEM. **p < 0.01, as determined by unpaired two-tailed Student’s *t* tests for 2 groups.

We also investigated if the secretome of senescent BMAds directly induces bone loss. BMAds isolated from AD and WT mice (Donors) were transplanted into the femoral bone marrow cavity of 3-month-old wild-type mice (Acceptors) (Figure 4D). Four weeks after transplantation, the bone phenotype of the Acceptors was examined. μCT analyses revealed that the Acceptors transplanted with BMAds isolated from AD mice (AD→WT), compared with those transplanted with an equal number of BMAds isolated from WT mice (WT→WT), had significant bone loss (Figure 4E), reduced BV/TV% (Figure 4F) and Tb.Th (Figure 4G), but unchanged Tb.N (Figure 4H) and Tb.Sp (Figure 4I). Thus, BMAds from AD mice alone are capable of inducing lower bone mass compared to those from WT mice.

### BMAds-secreted SAP promotes amyloid deposition, leading to increased osteoclast differentiation and decreased osteoblast differentiation

As SAP emerged as the most prominently released SASP factor in the antibody array screening (Figure 4B-C), we employed ELISAs to measure SAP concentrations in the CM collected from either AD mice (vs. WT mice) or old mice (vs. young mice) using the strategy illustrated in Figure 5A. As expected, BMAds from both AD mice (Figure 5B) and old mice (Figure 5C) showed elevated SAP secretion compared to their respective controls. To investigate whether increased SAP is associated with cellular senescence, we introduced either the combination of Dasatinib and Quercetin (D+Q), known for effectively eliminating senescent cells ^46^, or the JAK inhibitor (JAKi) ruxolitinib, recognized for inhibiting the SASP of senescent cells ^53,54^. In contrast to the significantly higher SAP concentration in the AD with vehicle treated group compared to the WT control group, both D+Q and JAKi treatment groups displayed reduced SAP concentrations, which were not significantly higher than that of WT group (Figure 5B). Consistently, SAP concentration in whole bone marrow was also significantly higher in AD mice (Figure S34A) and old mice (Figure S4B) compared to age-matched WT littermates and young mice, respectively. To further affirm that SAP is secreted by senescent BMAds, we conducted RNAscope analysis of bone tissue sections using a specific custom designed probe for *SAP* and a commercially available probe for *p21Cip*. While WT mice exhibited weak *p21Cip*^+^ signals on BMAds and undetectable *SAP*^+^ signal, AD mice displayed intense fluorescence signals for both *p21Cip* and *SAP*, and the colocalization of *p21Cip*^+^ and *SAP*^+^ signals was primarily in large, ring-like adipocytes (Figure 5D-E). Similarly, Old mice exhibited much stronger *p21Cip*^+^ and *SAP*^+^ signals in bone marrow compared to young mice (Figure 5F-G). Notably, almost all the *p21Cip/SAP* double-positive signals were localized at the large, ring-like structure, suggesting that SAP is primarily expressed in senescent BMAds.

**Figure 5.**
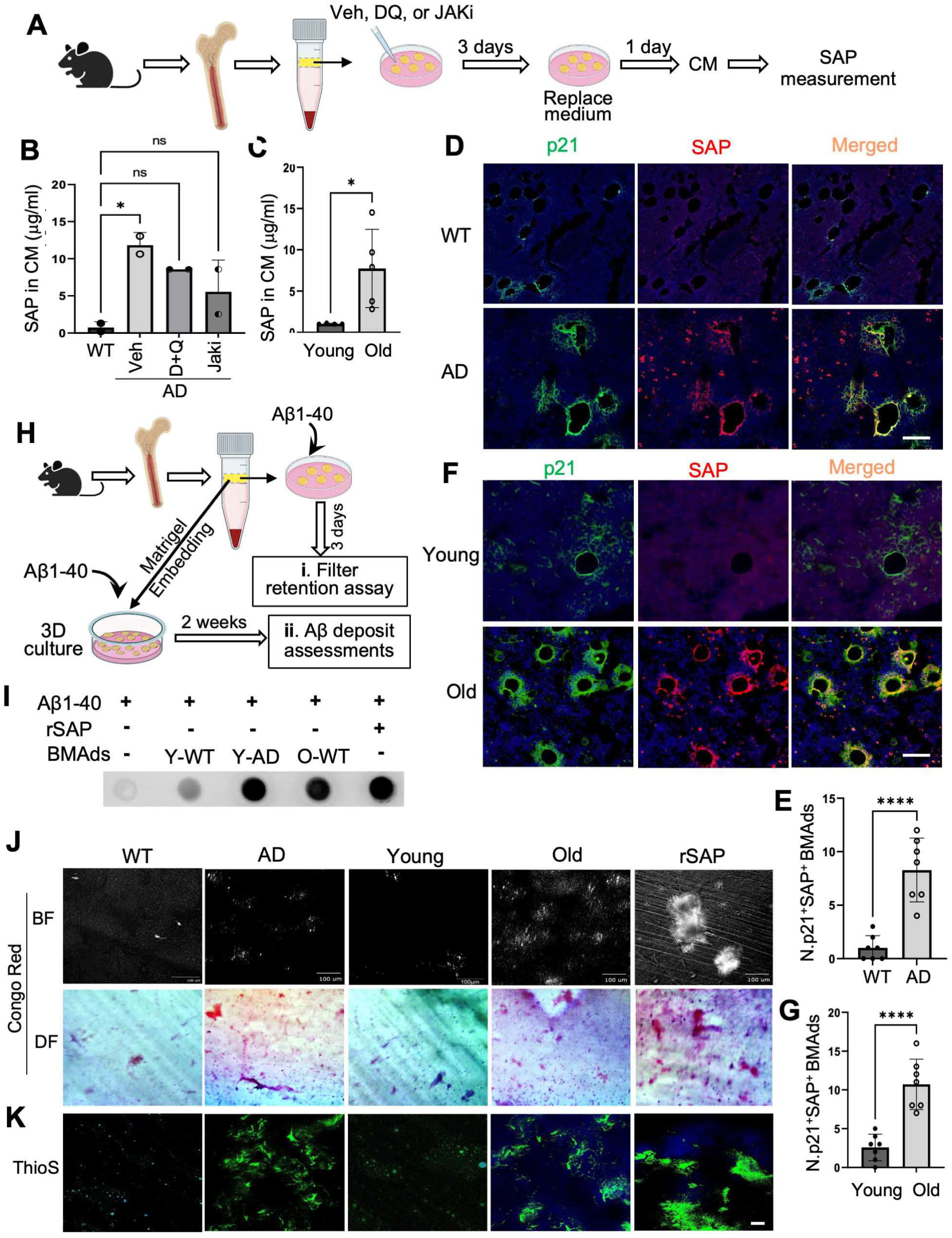
BMAd-secreted SAP drives amyloid deposition. (**A-C**) A schematic diagram illustrating the procedure of the SAP measurement in various conditioned media (CM) (**A**). BMAds isolated from either from 9-month-old *APP-PS1* mice (AD) and wild type littermates (WT) or from 24-month-old nature aging mice (Old and 4-month-old mice (Young) were cultured in aMEM with dasatinib and quercetin (DQ), JAK inhibitor (JAKi) ruxolitinib, or vehicle for 3 days. The secretion of SAP in the CMs were measured using ELISA. The concentrations of SAP in CM prepared from AD mice and WT mice were shown in (**B**) and from old and young mice were shown in (**C**) respectively. (**D-E**) Tibial bones were harvested from 9-month-old *APP-PS1* mice (AD) and their wild-type littermates (WT). Bone tissue sections were subjected to RNAscope assays using specific probes to *p21Cip* and *Apcs*. Representative images are shown in figure (**D**). Quantification of p21^+^SAP^+^ BMAds per mm^2^ tissue area were shown in (**E**). Scale bar = 100 pm. n=6-7. (**F-G**) Tibial bones were harvested from 24-month-old C57BL/6 mice (Old) compared to 4-month- old C57BL/6 mice (Young). Bone tissue sections were subjected to RNAscope assays using specific probes to *p21Cip* and *Apcs.* Representative images are shown in figure (**F**). Quantification of p21^+^SAP^+^ BMAds per mm^2^ tissue area were shown in (**G**). Scale bar = 100 pm. n=5-6. **(H-I)** A diagram illustrating the experimental procedure of the *in vitro* filter retention assays and *in vitro* Aβ deposit formation assay (H). BMAds isolated from different mouse strains were either cultured in adipogenic media and incubated for 3 days in the presence of freshly dissolved 60 pg/mL Aβ 1–40 followed by filter retention assays or cultured in a Matrigel-based 3D culture system. Filter retention assay results showing insoluble amyloid aggregates in each treatment group are shown in (**I**). Bright field (BF) or dark field (DF) polarizing microscopy and confocal Thio-S fluorescence images are shown in (**J**). Scale bar = 100 pm. n=6-7. Data are represented as mean ± SEM. *p < 0.05, ****p < 0.0001, as determined by unpaired two-tailed Student’s *t* tests for 2 groups.

To assess whether senescent BMAd-secreted SAP is sufficient to induce Aβ aggregation, we employed an *in vitro* BMAd culture system and a 3D BMAd culture platform, in which the formation of extracellular insoluable Aβ deposits can be detected by filter retention assays and amyloid staining approaches, respectively (Figure 5H). Filter retention assays of the *in vitro* culture (Figure 5Hii) revealed that the formation of insoluble Aβ aggregates was undetectable in Aβ1-40 peptide alone group (Figure 5I, the 1^st^ dot, serving as negative control). As a positive control, the incubation of recombinant human SAP (rSAP) with soluble Aβi -40 peptide led to a high intensity dot (Figure 5I, the last dot), suggesting the crucial role of SAP in promoting the formation of insoluble Aβ aggregates. Importantly, while the incubation of BMAds isolated from young WT mice with Aβ1-40 peptide resulted in the formation a weak insoluble Aβ aggregates (Figure 5I, the 2^nd^ dot), BMAds from AD mice and Old mice incubated with Aβ1-40 peptide formed dramatic more insoluble Aβ aggregates (Figure 5I, the 3^rd^ and 4^th^ dot). We also developed a 3D BMAd culture system (Figure 5Hii), in which BMAds isolated from mice were cultured in a 3D Matrigel matrix and incubated with Aβ1-40 peptide. Two weeks after Aβ1-40 incubation, aggregate formation was observed in BMAds from both AD mice and Old mice but not from age-matched WT mice and Young mice, as detected by the Congo red staining using both bright field (BF) and dark field (DF) polarizing microscopy (Figure 5J). ThioS staining reveals that the distribution and abundance of ThioS-positive plaques are consistent with that of Congo red-stained deposits in each treatment group (Figure 5K and Video 1). These findings indicate that BMAd-secreted SAP is sufficient to induce the accumulation of insoluble Aβ species. It is noteworthy that, serving as a positive control, direct mixing of rSAP with Aβ1-40 resulted in the largest amyloid deposit formation among all the groups (Figure 5J-K, last panel), further confirming the role of SAP in stabilizing insoluble Aβ fibrils.

To explore the impact of the Aβ aggregates on bone cell activities, we utilized an *in vitro* osteoclast-based cell culture system, treating isolated bone marrow monocytes/macrophages with M-CSF and RANKL to obtain mature osteoclasts. The combination of Aβ1-42 and rSAP largely increased osteoclastogenesis ability of the bone marrow Mo/Mac compared with vehicle treatment (Figure S5A and B). Additionally, we investigated the effect of Aβ aggregates on the activity of osteoblast lineage cells using bone marrow stromal cells (BMSCs), precursors of osteoblasts. Incubation with rSAP plus Aβ1-42 dramatically inhibited osteoblast differentiation ability, as detected by Alizerin Red staining of the cells at 21 days after the cells incubated with osteoblast differentiation medium (Figure S5C and D). The results suggest that Aβ aggregate formation induced by SAP promotes osteoclastogenesis but inhibits osteoblastogenesis.

### Depleting SAP using CPHPC (miridesap) effectively prevents the formation of amyloid deposits and rescues age-associated bone loss phenotype

To further determine the essential role of SAP in promoting the formation of bone amyloid deposits and associated bone impairment, we utilized a well-established bivalent small molecule drug called CPHPC (miridesap). This compound efficiently binds to SAP, leading to the dissociation of SAP from bound amyloids and subsequently depletion of SAP levels in the body ^24,55–61^. After administering CPHPC to 18-month-old mice for 6 weeks, the concentrations of SAP were markedly reduced in both bone marrow and serum (Figure 6A-B), validating the effectiveness of the drug in depleting SAP. Analyses of bone phenotype revealed significantly higher bone mass of distal femur in CPHPC-treated female mice relative to the control mice with normal drinking water (Figure 6C). Quantification of the parameters in the trabecular bone showed significantly increased BV/TV%, Tb.Th, unchanged Tb.N, but decreased Tb.Sp, in the CPHPC treatment group relative to the control group (Figure 6D-G). Consistent results were obtained from proximal tibia in female mice (Figure S6A-D) as well as distal femur (Figure S6E-H) and proximal tibia in male mice (Figure S6I-L). CPHPC-treated mice also demonstrated increases in cortical bone area (Ct.Ar) and cortical thickness (Ct.Th) compared with the control mice (Figure 6H-J). We then conducted histological analysis of the bone tissue sections. Consistent with Figure 2K, abundant H31L21^+^ and ThioS^+^ amyloid deposits were detected in bone marrow of aged mice, whereas these amyloid deposits were almost eliminated in CPHPC-treated mice (Figure 6K-M). We observed greatly reduced TRAP^+^ osteoclasts with much lower surface per bone surface (Oc.S/B.S.) in both trabecular bone (Figure 6N and O) and cortical bone (Figure 6N and P) in CPHPC-treated mice relative to vehicle treated mice. Together, the *in vivo* data provide evidence to support the role of SAP in inducing bone amyloidosis, which leads to bone loss.

**Figure 6.**
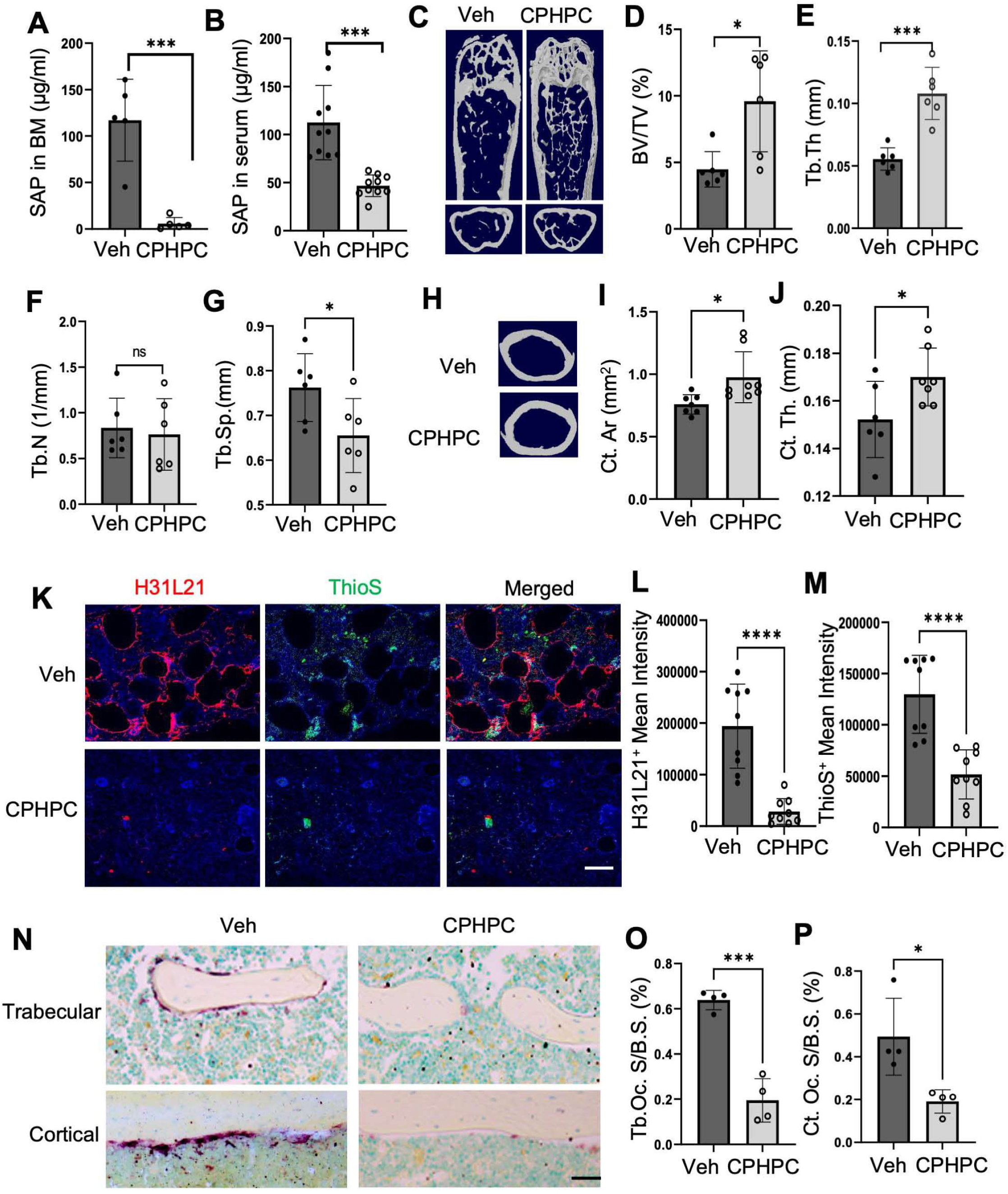
Depleting SAP by CPHPC (miridesap) alleviates amyloid deposits and improves bone phenotype in aged mice. Eighteen-month-old male and female C57BL/6 mice were treated with CPHPC at 5mg/ml through drinking water for 42 days. Regular drinking water was used as a control. (**A-B**) The concentrations of SAP in serum (**A**) and bone marrow (**B**) were measured by ELISA. (**C-J**) Representative μCT images of the trabecular area in the distal femur of female mice are shown in (**C**) with quantitative analyses of trabecular bone volume (BV/TV) (**D**), trabecular thickness (Tb. Th) (**E**), trabecular number (Tb. N) (**F**), and trabecular separation (Tb. Sp) (**G**). n=6. Representative μCT images of the cortical area of the femur are shown in (**H)** with quantitative analyses of cortical area (Ct. Ar) **(I)** and cortical thickness (Ct. Th) **(J)**. n=6. (**K-M)** Aβ deposits were detected by ThioflavinS (ThioS) staining, Congo red staining and immunofluorescence staining of bone tissue sections with antibody against H31L21. Representative images were shown in (**K**). Quantifications of the ThioS+ Mean mean intensity and H31L21+ mean intensity were shown in (**L**) and (**M**), respectively. Scale bar = 100 pm. n=9. (**N-P**) Representative TRAP staining images in femoral trabecular and cortical regions (**N**) with quantification of TRAP+ osteoclast surface per bone surface (Oc.S./B.S.) in trabecular area (**O**) and cortical area (**P**). n = 4 mice for each group. Scale bar = 50 pm. Data are represented as mean ± SEM. *p < 0.05, ***p < 0.001, ****p < 0.0001, as determined by unpaired two-tailed Student’s *t* tests for 2 groups.

## DISCUSSION

While age-related amyloid has been observed in several non-brain tissue sites, almost nothing is known about the development of skeletal amyloidosis and its potential contribution to age-associated skeletal disorders. Our study unveils that senescent BMAds-secreted SAP/PTX2 is a major contributor to bone marrow amyloidosis and subsequent bone impairment. BMAds undergo substantial expansion and display a senescence phenotype in both AD mice and nature aging mice. These senescent BMAds acquire a secretory phenotype, leading to the secretion of elevated levels of SAP/PTX2, which in turn stabilizes amyloid fibrils and promotes amyloidogenesis around BMAds. The presence of amyloid fibrils triggers osteoclast differentiation and impedes osteoblastogenesis, ultimately resulting in bone loss (Figure 7). These findings provide evidence that strategies aimed at antagonizing BMAd senescence or depleting/reducing SAP/PTX2 may represent therapeutic opportunities for improving bone health in the elderly.

**Figure 7.**
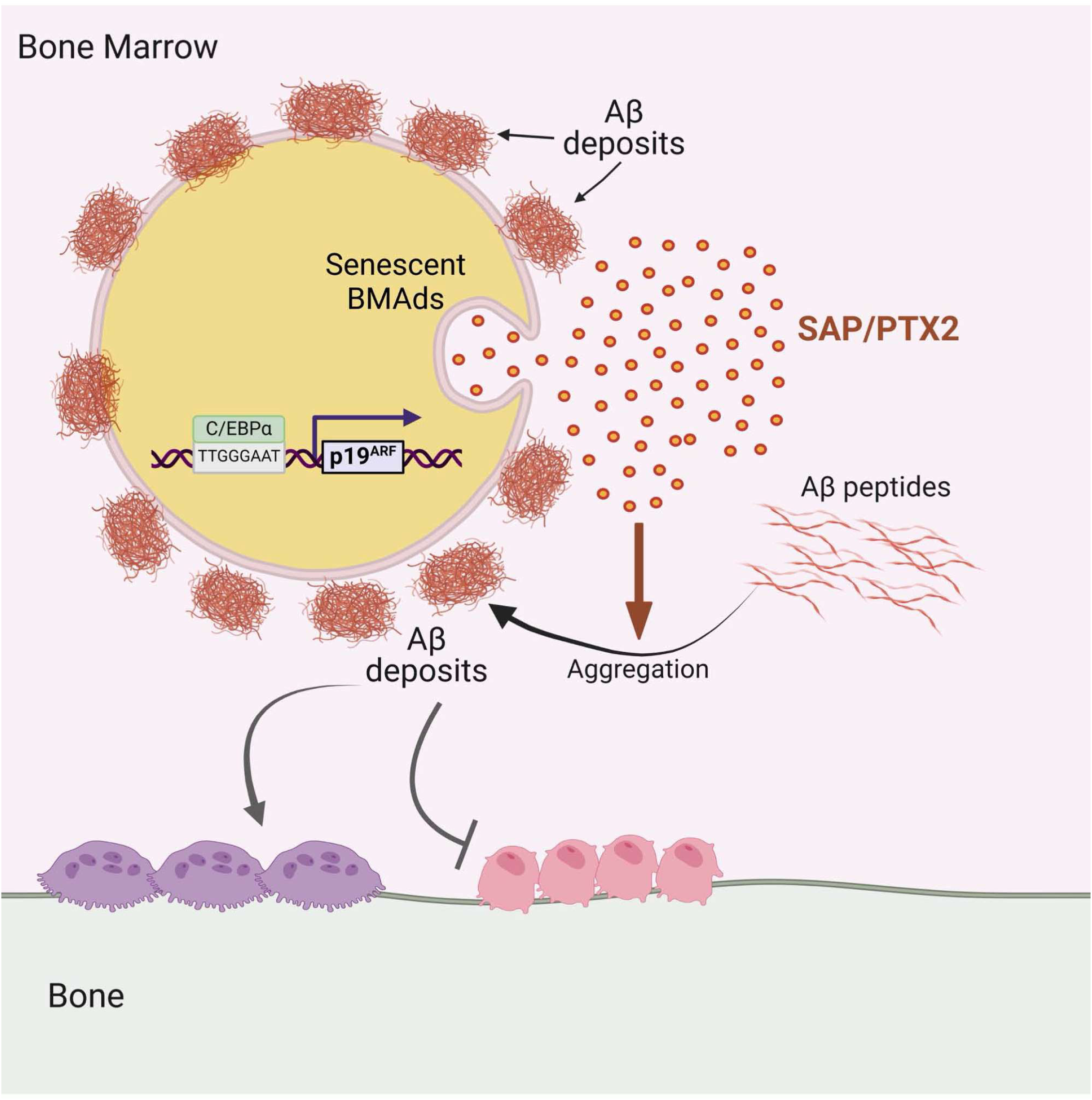
Diagram illustrating the role of senescent BMAd-secreted SAP/PTX2 in the formation of Aβ deposits in their vicinity.

Our data reveals an accumulation of senescent BMAds in both AD mice and aged mice, characterized by increased mRNA and protein expression of a panel of senescence markers such as the DNA damage marker γH2AX, and tumor suppressors p16Ink4a, p19Arf, and p21Cip as detected by either RNAscope or immunofluorescence staining. Despite sharing largely overlapping DNA sequences, the *Ink4a/Arf* locus encodes two completely distinct proteins (p16Ink4a and p19Arf). Previous studies revealed that PPARγ is a key transcription factor mediating *p16Ink4a* gene activation ^48^ and BMAd senescence ^42^. Here we identify a distinct molecular mechanism underlying the activation of *p19Arf* gene, the role of which in mediating cellular senescence has been overshadowned by *p16Ink4a* for many years. Intriguingly, ChIP assays reveal that transcription factor CEBPα strongly binds to all four regions in the promoter of *p19Arf* in bone marrow cells from both AD mice and aged mice. However, the bindings of CEBPα to the *p16Ink4a* promoter region were not detected in either WT or AD mice, although *p16Ink4a* promoter also contains four putative CEBPα binding sites. The data suggests that while *p16Ink4a* and *p19Arf* are in the same locus and share overlapping coding regions, they are independently activated in BMAds via distinct mechanisms during aging or in AD conditions. It is of interest to determine whether *p16Ink4a* and *p19Arf* are activated simultaneously or sequentially in the progression of aging or AD. It is also important to examine whether these two closely-related senescence marker genes are differentially activated in the senescence of other cell types. Another notable feature of senescent BMAds is their acquisition of a SASP. In the cytokine/adipokine array assays, 17 out of 111 cytokines/adipokines were elevated in BMAds isolated from AD mice compared to their control compartments. It is noteworthy that the majority of the elevated factors in this context differ from the SASP factors previously identified in glucocorticoid-induced senescent BMAds ^42^. These results support the consensus recognition from numerous other studies that the composition of SASP released from senescent cells can be highly specific and dynamic, contingent upon the senescence inducer and the duration of senescence ^46,62–67^. Nevertheless, the findings from the present study and our recent work on glucocorticoid-induce BMAds collectively suggest that BMAds are more susceptible or rapidly acquire a senescence phenotype compared to other cell types in the bone/bone marrow, likely because the master regulators of adipogenesis such as PPARγ and CEBPα also functions as senescence gene activators.

Perhaps the most striking observation of the current work was the presence of Aβ fibril accumulation around senescent BMAds. We observed plaque-like Aβ deposits in the bone marrow of AD mice using ThioS staining and APP immunostaining. In aged wild-type mice, smaller Aβ deposits are either diffuse in the bone marrow or localized around senescent BMAds, forming unique large, ring-like structure. Although it is commonly understood that wild-type mice and rats typically do not develop Aβ plaques like humans with AD ^2,68,69^, the expressions of amyloid precursor protein (APP) and soluble Aβ were previously detected in aged WT bone marrow cells ^70^. Moreover, extensive Aβ deposits have been identified in the brains of SAMP8 mice ^71–75^, an accelerated aging mouse strain lacking human *APP* transgene. Our study provides the first evidence that insoluble Aβ fibrils aggregate in the bone marrow of aged mice, albeit not in the form of large, dense Alzheimer’s-like plaques. The distinctive peri-BMAd feature of bone marrow Aβ deposits serves as a hallmark characteristic of skeletal aging.

A fundamental question arises regarding the mechanism underling the accumulation of insoluble Aβ fibrils in the bone marrow with age. Given our observation that amyloid aggregates predominantly around BMAds, it is logical to posit that Aβ deposition is associated with the aging phenotypic changes of this cell type. Indeed, our *in vitro* and *in vivo* assays reveal that senescent BMAds in AD mice and aged mice secrete significantly higher amounts of SAP/PTX2, thereby promoting the formation of insoluble Aβ aggregates. Previous studies have shown that SAP/PTX2 is primarily produced in the liver in physiological conditions in humans ^76–78^. Our observation from RNAscope assay, showing that SAP/PTX2 largely colocalizes with the senescence marker gene *p21Cip* in BMAds, suggests that SAP/PTX2 is produced by senescent BMAds. Supporting this notion, SAP/PTX2 levels were markedly elevated in the conditioned media prepared from BMAds isolated from both AD mice and nature aging mice compared to their control counterparts, and this elevation was mitigated by treating the cells with senolytic drugs. Furthermore, SAP/PTX2 concentrations in bone marrow of aged (vs. young) mice and AD mice (vs. WT littermates) were significantly elevated. These findings collectively suggest that BMAds are a primary source of SAP/PTX2 in the bone marrow microenvironment in mice during aging or under AD conditions. It has been well-established that SAP is universally present in all human amyloid deposits ^61,79^. The binding of SAP stabilizes amyloid fibrils ^80^ and promotes their formation in the brain ^81,82^, thereby contributing to the pathogenesis of AD and cerebral amyloid angiopathy ^83,84^. Our results from filter retention assays and the 3D amyloid deposition assays demonstrate that senescent BMAds-secreted SAP/PTX2 efficiently triggers the formation of insoluble amyloid fibrils from soluble Aβ peptide, providing evidence for the role of SAP in stabilizing amyloid fibrils in peripheral tissue.

The discovery of SAP-induced amyloid deposits in the bone marrow may pave the way for a new understanding of age-associated bone loss. Our *in vitro* assays demonstrate that combined SAP and Aβ peptide treatment increased osteoclastogenesis of bone marrow monocytes/macrophages and inhibited osteoblastogenesis of bone marrow mesenchymal stromal cells, suggesting that SAP-triggered Aβ deposition may regulate the function of both osteoclast and osteoblast lineage cells. This likely explains why AD mice with accumulated Aβ deposition in the bone marrow develop a similar low bone mass phenotype as observed in aged mice. Our data demonstrated that CPHPC (miridesap), developed by Pepys’ team at the University College London ^85–87^, efficiently reduces SAP concentration in serum (58.5% reduction) and bone marrow (95% reduction) in aged mice. Intriguingly, both aged male and female mice exhibit abolished bone marrow Aβ deposits with markedly increased bone mass after 6 weeks of CPHPC treatment compared to vehicle-treated mice. Moreover, CPHPC treatment reduces osteoclast number and increases osteoprogenitor cells, indicating the impact of depleting SAP on both osteoclast and osteoblast activities. This *in vivo* result, combined with the *in vitro* data, suggests a significant impact of SAP-induced bone marrow Aβ deposits on bone cell activities. It is important to note that CPHPC was developed based on the human SAP sequence, and its SAP-depletion effect has primarily been tested in humans. However, consistent with our mouse result, several recent studies have reported that this drug is also effective in depleting SAP in mice in different disease models ^55,57^. Further validation of the bone-improvement effect of CPHPC in other pathological settings, such as glucocorticoid- or postmenopausal-induced bone loss, will provide a better understanding on the role of SAP in the pathogenesis of skeletal disorders.

## METHOD DETAILS

### Animals and treatment

#### Mouse lines

In this study, we mainly used Male and female mice at the age of 4, 9, 18, 22, and 24 months. Mouse strains used were: C57BL/6J mice obtained from National Institute on Aging; *APPswe/PS1ΔE9 (APP-PS1)* obtained from Jackson Lab (MMRRC Strain #034832-JAX) and maintained as double hemizygotes by crossing with wild-type individuals on a C57BL/6 background. Tail biopsies samples were used to isolate DNA to confirm the genotype of the strain using previously established primers by Polymerase chain reaction. ^88–90^. To examine the effects of CPHPC on age-associated bone changes, male and female mice aged 18 months were treated with CPHPC (miridesap) at a concentration of 5 mg/ml in drinking water, which is an established dosage for SAP depletion from the body ^55–58,60,61,79^. Control mice were treated with the same amount of water without the drug. On day 42nd, mice were sacrificed, and serum and bone marrow SAP concentrations were measured using ELISA. The bone phenotype was assessed by micro-CT and histological analyses.

Mice were kept under standard conditions with a 12-hour light-dark cycle in a temperature-controlled room and provided with food and water ad libitum. They were housed at The Johns Hopkins University School of Medicine’s animal facility in Baltimore, Maryland, following protocols MO21M37 and MO24M35 approved by the Institutional Animal Care and Use Committee. All procedures adhered to the guidelines of the Institutional Animal Care and Use Committee.

### μCT analyses of bone phenotype and bone marrow adiposity

Femoral and tibial bones were dissected from mice, free of soft tissue, fixed overnight in 10% formalin at 4°C, and subjected to high-resolution μCT imaging using a Skyscan 1172 scanner (Bruker MicroCT in Kontich, Belgium). The scanning parameters were set at 65 kV, 153 μA, with a resolution of 9.0 μm/pixel. Bruker MicroCT software tools, including NRecon for image reconstruction (version 1.6), CTAn for data analysis (version 1.9), and CTVol for 3-dimensional model visualization (version 2.0) were used to analyze parameters of the trabecular bone in the metaphysis and mid-diaphyseal cortical bone. Trabecular bone parameters, specifically trabecular bone volume fraction (BV/TV%), trabecular thickness (Tb.Th), trabecular number (Tb.N), and trabecular separation (Tb.Sp) as well as cortical bone parameters, including cortical area (Ct. Ar) and cortical thickness (Ct. Th), were quantified as previously described ^91,92^.

Bone marrow adiposity of mice was measured following established procedures ^42,93^. Briefly, femurs harvested from mice were fixed in 4% phosphate-buffered paraformaldehyde for 48 hours, and decalcified for 2 weeks in 10% EDTA at 4°C. The distal parts of femurs were incubated in 2% aqueous OsO4 (MilliporeSigma) for 2 hours in the fume hood. The femurs were rinsed in PBS and then scanned using a high-resolution μCT scanner (SkyScan 1172, Bruker MicroCT) at 6-pm resolution using 45 peak kV (kVp) and 177 μA. Quantification of adipocyte volume/tissue volume (AV/TV%) was conducted.

### Tissue section preparation, histological analysis, and immunostaining

The femur and tibia bones were dissected and fixed in 4% paraformaldehyde overnight, followed by a 21-day decalcification in 0.5M EDTA (pH 7.4). Afterward, the samples were dehydrated in a solution of 20% sucrose and 2% polyvinylpyrrolidone for 24 hours and embedded in OCT. Ten-micrometer-thick coronal sections of the femurs were prepared for SA-βGal staining. Senescent cells were identified using a senescence pGal staining kit (Cell Signaling Technology, Danvers, MA) following the manufacturer’s instructions. For Oil Red O staining after SA-βGal staining, the slides were washed in PBS three times, immersed in 100% propylene glycol for 2 minutes, and then stained in a filtered 0.5% Oil Red O solution in propylene glycol for 30 minutes. Subsequently, the slides were placed in an 85% propylene glycol solution for 1 minute, rinsed in PBS three times, mounted, and sealed with nail polish. Representative images were captured using a BX51 microscope (Olympus, Tokyo, Japan).

Twenty μm-thick coronal sections of the femurs were obtained for immunofluorescence staining. Bone sections were blocked in PBS with 3% BSA for 1 hour and then stained overnight (>8 hours) with individual primary antibodies *p19Arf* (Novus NB200-111, 1:100), γH2AX (Cell signaling, 20E3, 1:200), Perilipin (Cell signaling, 9349, 1:200), Perilipin (Sigma, P1873, 1:500), and H31L21 (Thermo Fisher, Scientific 700254, 1:100). Fluorescence-conjugated secondary antibodies (Jackson ImmunoResearch, 1:200) were used in immunofluorescence procedures to detect fluorescent signals. Slides were then mounted with anti-fade prolong gold (Invitrogen) mounting medium and sealed with nail polish, after which representative images were acquired using a Zeiss LSM780 confocal microscope or an Olympus BX51 microscope. TRAP staining was performed in selected slides from each sample using a commercially available kit (Sigma-Aldrich, 387A) according to the manufacturer’s instructions. Images were captured using a BX51 microscope (Olympus, Tokyo, Japan), and quantitative analysis was performed using Image J software (v.1.54f, National Institutes of Health, Bethesda, MD, USA). For ThioS staining, tissue sections were incubated with 1% ThioS (Sigma-Aldrich, T1892) in distilled water for 8-10 mins, followed by rinsing with 50% ethanol twice, slides were then mounted with anti-fade prolong gold (Invitrogen) mounting medium before imaging.

### RNAscope

RNAscope in situ hybridization (ISH) was conducted following the guidelines provided by Advanced Cell Diagnostics (ACD; Hayward, CA). Probes specific to *Cdkn1a* (p21) (Cat No. 408551), *Cdkn2a* (*p19Arf*) (Cat No. 411011), *Cdkn2a* (p16) (Cat No. 447491), *H2AX* (Cat No. 446191), and custom designed SAP/PTX2 encoding gene *Apcs* (Cat No. 1304211-C3) were employed according to the manufacturer’s protocol. Briefly, frozen tissue slides were fixed in 10% formalin and underwent series dehydration. Protease III reagent was applied to each section. Subsequently, a mixture of Channel 1, Channel 2, and Channel 3 probes (50:1:1 dilution, per ACD’s instructions based on stock concentrations) was pipetted onto each section until fully submerged. The humidity control tray, with slides, was then incubated in a HybEZ oven (ACD) for 2 hours at 40°C. All experiments featured a target of interest in Channel 1 in conjunction with a probe in Channel 2. Following probe incubation, the slides underwent two washes in 1X RNAscope wash buffer and were returned to the oven for 30 minutes after submersion in AMP-1 reagent. Washes and amplification were repeated using AMP-2 and AMP-3 reagents with 15- minute and 30-minute incubation periods, respectively. Slides were then incubated with fluorophores for respective channels and with DAPI.

### Isolation of BMAds from mice

Adipocytes were directly extracted from the bone marrow of mice following established procedures ^94,95^. Briefly, femurs and tibias were harvested from the mice, and the bone ends were trimmed. These bones were then placed in a small microcentrifuge tube (0.6 mL) with an open bottom, which was subsequently inserted into a larger microcentrifuge tube (1.5 mL). Fresh bone marrow was separated out by quick centrifuge. Red blood cells were lysed using RBC lysing buffer (Sigma). Lysed bone marrow was centrifuged at 3000 rpm, for 5 minutes and floating adipocytes were collected from the top layer and washed 3 times with PBS.

### Cell Transplantation

To assess the effect of senescent BMAds on regulating bone remodeling, BMAds isolated from *APP-PS1* mice (Donors) were transplanted into healthy wild-type C57BL/6 mice via intra-femoral transplantation. Briefly, the recipient mice (Acceptors) were anesthetized, and a longitudinal skin incision was made across the front of the right knee to expose the patellar tendon A 27G needle was carefully inserted through the patellar tendon, between the femur condyles, and advanced slowly in a twisting motion through the bone surface to a depth of 2-4 mm. Successful penetration was confirmed by a slow but consistent bleeding response. Cell suspensions of BMAds (3*10^3^) in 20 μl PBS were gradually injected into the medullary space of the femur. The resulting hole was promptly sealed with bone wax, and the skin was sutured. After four weeks, mice were sacrificed, and the bone phenotype was subjected to analysis.

### Proteomic profiling in conditioned medium (CM) collected from BMAd culture

BMAds were isolated from the long bones and cultured in a growth factor-free medium for 72 hours to collect CMs, which were then stored at -80°C for subsequent experiments. Cytokine levels were assessed using a mouse XL cytokine array kit (R&D, ARY028) and a Mouse Adipokine Array Kit (R&D, ARY013) through a proteome profiler array. The measurement of relative cytokines and chemokines levels, as well as the array procedure and data analysis, followed the manufacturer’s instructions.

### qRT-PCR

Total RNA for quantitative reverse transcription polymerase chain reaction (qRT-PCR) was extracted from total bone marrow cells utilizing the RNeasy Mini Kit (QIAGEN), following the manufacturer’s protocol. Complementary DNA (cDNA) was synthesized with random primers using the SuperScript First-Strand Synthesis System (Invitrogen) and analyzed with SYBR GreenMaster Mix (QIAGEN) in a thermal cycler employing two sets of primers specific for each targeted gene. The expression of the target genes was normalized to glyceraldehyde 3-phosphate dehydrogenase (GAPDH) messenger RNA, and the relative gene expression was determined using the 2–ΔΔCT method. The primer sequences employed for qRT-PCR were as follows: *p19Arf* (Forward -5’-CGCAGGTTCTTGGTCACTGT-3’) and Reverse 5’- TGTTCACGAAAGCCAGAGCG-3’). *p16Ink4a* (Forward-5’- AACTCTTTCGGTCGTACCCC -3’) and Reverse-5’- GCGTGCTTGAGCTGAAGCTA -3’). *P21Cip* (Forward-5’- GCAGATCCACAGCGATATCCA-3’) and (Reverse-5’- TTTCGGCCCTGAGATGTTCC-3’)

### Bone marrow stromal cell culture and osteogenic differentiation

Bone marrow stromal cells (BMSCs) were isolated from tibiae and femurs of 6 weeks old C57BL/6J mice using previously established protocols ^93,96^. Briefly, tibiae and femurs were cut and spun by centrifuge, cells were then resuspended in complete a-MEM with 1% penicillin/streptomycin and 10% FBS. The first passage cells were used for plating and further experiment. When cells reached 75–85% confluency, osteoblastogenesis of BMSCs were initiated using osteogenic induction media (Stemcell tech., Cat no. 05504). Medium was changed every two days. To assess mineralization, Alizarin Red staining was performed on Day 21 after differentiation using a commercially available kit (Millipore Sigma, USA, Cat no. 2003999), and the manufacturer’s instructions were followed for the procedure^97^. Cells were then washed a couple of times with water before being visualized under a BX51 microscope (Olympus, Tokyo, Japan). Cells were then stored at -20 °C until the photometric analysis.

### *In vitro* osteoclast culture

*In vitro* osteoclastogenesis assays were performed using previously established protocols ^96,98^. Briefly, monocytes/macrophages were isolated from 4-week-old C57BL/6J mice by flushing cells from the bone marrow of tibiae and femora. The flushed bone marrow cells were cultured with ot- MEM containing 15% FBS, 100U/mL streptomycin sulfate and 100U/mL penicillin. The cells were then incubated with osteoclastogenesis medium, containing aMEM with 10% FBS, 1% penicillin-streptomycin solution, 30 ng/mL M-CSF and 100 ng/mL RANKL (Amizona Scientific LLC, AM10004-500) for 5 days to induce the formation of mature osteoclasts.

### ELISA analysis of Aβ peptides and SAP/PTX2 concentrations

The Amyloid beta 42/40 Mouse ELISA Kit (KMB3441/KMB3481, Thermo Fisher Scientific) was then used to determine the concentrations of amyloid beta 42/40 in mouse bone marrow. The Mouse Pentraxin 2/SAP Quantikine ELISA Kit (MPTX20, R&D Systems) was utilized to quantify the SAP/PTX2 concentration in mouse bone marrow and in serum. The assessments were conducted using a spectrophotometer (A51119600DPC, Thermo Fisher Scientific).

### ChlP-qPCR

ChlP-qPCR was performed using the Agarose Magnetic ChIP kit (PI26157, Thermo Fisher Scientific) following the manufacturer’s instructions. Immunoprecipitation was performed using a valid CEBPα chip antibody (Cat #PA5-77911, Thermo Fisher Scientific). The RNA polymerase II antibody as a positive control was provided in the kit. The primers used for PCR were as follows: mouse- CEBPα _Fwd-5’ATTCCAGGTAGGGTAGCCCG-3’ and Rev- 5’TGTGACTTTGCAGTGGAACCT-3’; the GAPDH primer was provided in the kit.

### Detection of amyloid plaque formation in an *in vitro* BMAd Culture by filter retention assays

*In vitro* amyloid plaque formation in cell culture system was performed as previously described ^99^ with modification. Briefly, BMAds isolated from mice were cultured in 6-well plates, and freshly dissolved Aβ1-40 peptide at a final concentration of 60 pg/ml was supplemented in the culture medium. Medium will be replaced every two days. Two week later, cells in 6-well plates were thoroughly washed with PBS and then lysed in 150 pl RIPA buffer containing a protease inhibitor mix for 30 minutes on a shaking device at 4 °C. After adding 150 pl of water and 100 pl of 4% sodium dodecyl sulfate, 100 mM dithiothreitol, the solution was boiled for 2 x 7 min at 98°C and filtered through a 0.2 pm cellulose acetate membrane (Bio-Rad) with a 96-well vacuum Bio-Dot Apparatus (Bio-Rad; cat# 1706545), followed by three washing steps with lx TBST. The membrane was blocked for 1 h in 3% gelatin-TBS. Aggregates were detected by incubation with anti-Aβ (1-16) primary antibody (BioLegend; 1:200 in blocking buffer) overnight, followed by incubation with AP conjugated anti-mouse secondary antibody (1:15,000 in blocking buffer; Sigma) for another hour. Blots were developed with LumiPhos reagent according to the manufacturer’s instructions (Pierce), and the signals were scanned and quantitatively analyzed with iBright Imaging Systems (Thermo Fisher Scientific - US).

### Detection of amyloid plaque formation in an *ex vivo* 3D BMAd Culture platform

3D cultures were established as previously described ^100^. Briefly, Matrigel (Cat # DLW356231; Sigma-Aldrich) was added to the cell suspension on ice in a 1:1 (vol/vol) dilution. The pipette tips were chilled by pipetting cold differentiation medium back and forth before transferring the Matrigel. The 1:1 Matrigel/cell mixture was further diluted by adding five volumes of cold MesenCult™ Adipogenic Differentiation Kit (Mouse) (Cat # 05507; stemcell tech, USA) (1:11 dilution final) and vortexed for 30 seconds. 200 pl of the Matrigel/cell suspension mixture was plated per well of an eight-well chambered cover-glass slide using prechilled pipettes, and the plates were incubated at 37 °C in a CO2 incubator overnight. The next day, two drops (200 pl) of prewarmed adipocyte differentiation medium were added to each well of the eight-well chambered cover-glass slides. Half of the medium volume was changed every 3-4 days, initially changing the medium every 2 days depending on the medium color. Plaque formation was induced by supplementing the culture medium with freshly dissolved Aβ1-40 peptide at concentration of 60 pg/mL. Freshly dissolved Aβ was replenished each time the culture medium was replaced.

To detect the amyloid plaque formation, Congo red green birefringence was carried out as described previously ^99^. Briefly, cells grown on 8-well chambered cover-glass slides were fixed in ice-cold methanol for 10 minutes, and stained for 45 minutes with filtered Congo red solution [80% ethanol, 3% NaCl, 0.6% Congo red]. The cells were then washed three times with water and incubated for 2 minutes with Mayer’s Hematoxylin Solution, followed by one wash with 70% ethanol and three washes with water. Subsequently, the cells were mounted and examined with The Olympus 1X83 inverted microscope (JHU SOM Microscope facility). ThioS staining was also used for the detection of amyloid plaque formation in the 3D cultures. Samples were fixed, permeabilized, and blocked, and incubated with 1% Thio S in distilled water for 8-10 mins, followed by rinsing with 50% ethanol twice, slides were then mounted with anti-fade prolong gold (Invitrogen) mounting medium before imaging.

### Statistical analysis

The data are expressed as means ± standard errors of the mean. Unpaired, two-tailed Student /-tests were employed for comparisons between two groups. For multiple comparisons, one-way analysis of variance with Bonferroni post hoc test was utilized. All data exhibited normal distribution and had comparable variation between groups. Statistical analysis was conducted using g GraphPad Prism 8 statistical software (GraphPad Software Inc., La Jolla, CA). A significance threshold of p < 0.05 was applied. Representative images of bones or cells were chosen from a minimum of three independent experiments with consistent results unless specified otherwise in the figure legend.

## AUTHOR CONTRIBUTIONS

S.K. and M.W. designed the experiments, analyzed results, and wrote the manuscript; S.K. carried out most of the experiments; K.S., J.W., and M.S.B. helped with some experiments; M.W. supervised the experiments; X.C., and P.W. proofread the manuscript.

## ACKNOWLEDGEMENT

We acknowledge the assistance of the Johns Hopkins Ross Flow Cytometry Core Facility (supported by NIH shared-instrument grant) and School of Medicine Microscope Facility. This work was supported by the National Institutes of Health grant R01AG068226 and R01AG072090 to M.W., and P01AG066603 to X.C.

## COMPETING INTERESTS

The authors declare no competing financial interests.

## Supplemental Text and Figures

**Figure S1.**
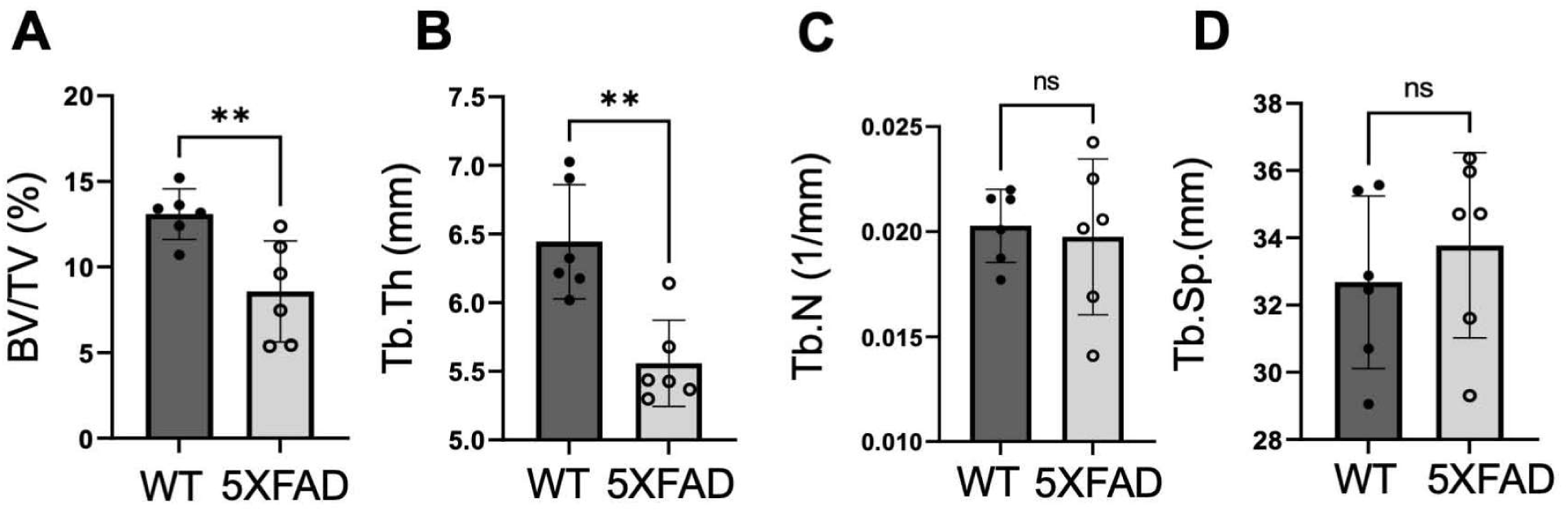
5XFAD mice exhibit low bone mass phenotype. Femoral bones were harvested from 13-month-old male 5xFAD mice and their WT littermates. 5xFAD mice express human APP and PSEN1 transgenes with a total of five AD-linked mutations: the Swedish (K670N/M671L), Florida (1716V), and London (V717I) mutations in APP, and the M146L and L286V mutations in PSEN1. Mico-CT analyses of the trabecular area in the distal femur were performed. Quantifications of trabecular bone volume (BV/TV%) **(A),** trabecular thickness (Tb. Th) **(B)**, trabecular number (Tb. N) **(C)**, and trabecular separation (Tb. Sp) **(D)** are shown. n=6 mice. Data are represented as mean ± SEM. **p < 0.01, as determined by unpaired two-tailed Student’s *t* tests for 2 groups

**Figure S2.**
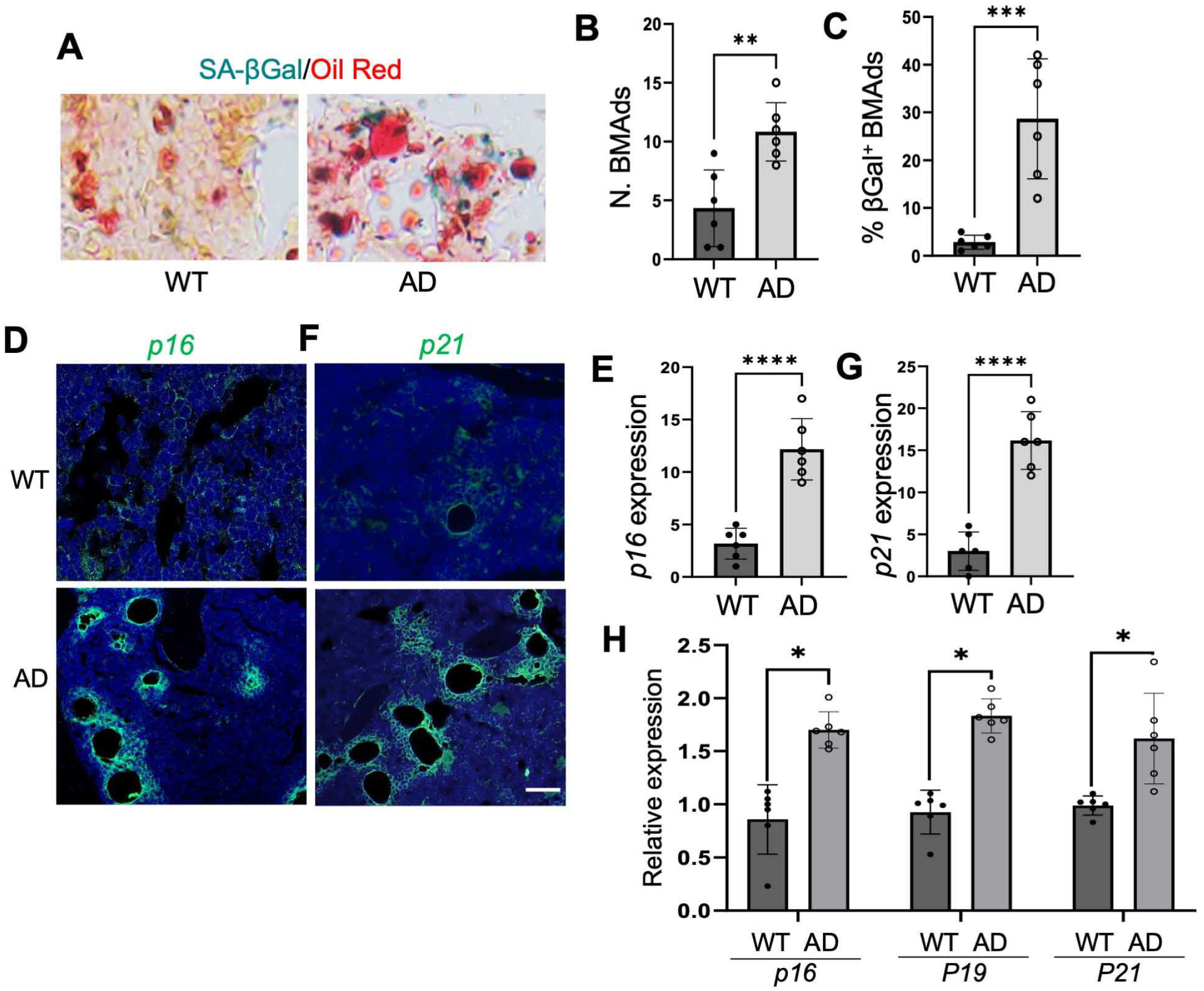
BMAds in AD mice undergo cellular senescence. Nine-month-old APP-PS1 mice (AD) and wild type littermates (WT) were sacrificed, and bone tissue sections were prepared. (A-C) Representative images of SA-pGal staining and Oil Red 0 staining are shown in (A). Quantifications of the total numbers of Oil Red O+ adipocytes per mm2 tissue area (N. BMAds) and the percentages of SA-pGal-expressing BMAds (% pGal + BMAds) are shown in (B) and (C), respectively. Scale bar, 50 pM. n=6 mice. (D-G) Bone tissue sections were subjected to RNAscope assays using specific probes to pl6Ink4a and p21Cip1. Representative images are shown in (D) and (F), respectively. Quantifications of pl9ARF + mean intensity and H2AX + mean intensity are shown in (E) and (G), respectively. Scale bar =100 pm. n=6. (H) Bone marrow cells isolated from AD and WT mice were subjected to qRT-PCR for detection of pl6Ink4a, pl9Arf, and p21Cip mRNA expression. Data are represented as mean ± S.E.M. *p < 0.05, **p < 0.01, ***p < 0.001, as determined by unpaired two-tailed Student’s t test for 2 groups and one­way ANOVA for 3 or more groups.

**Figure S3.**
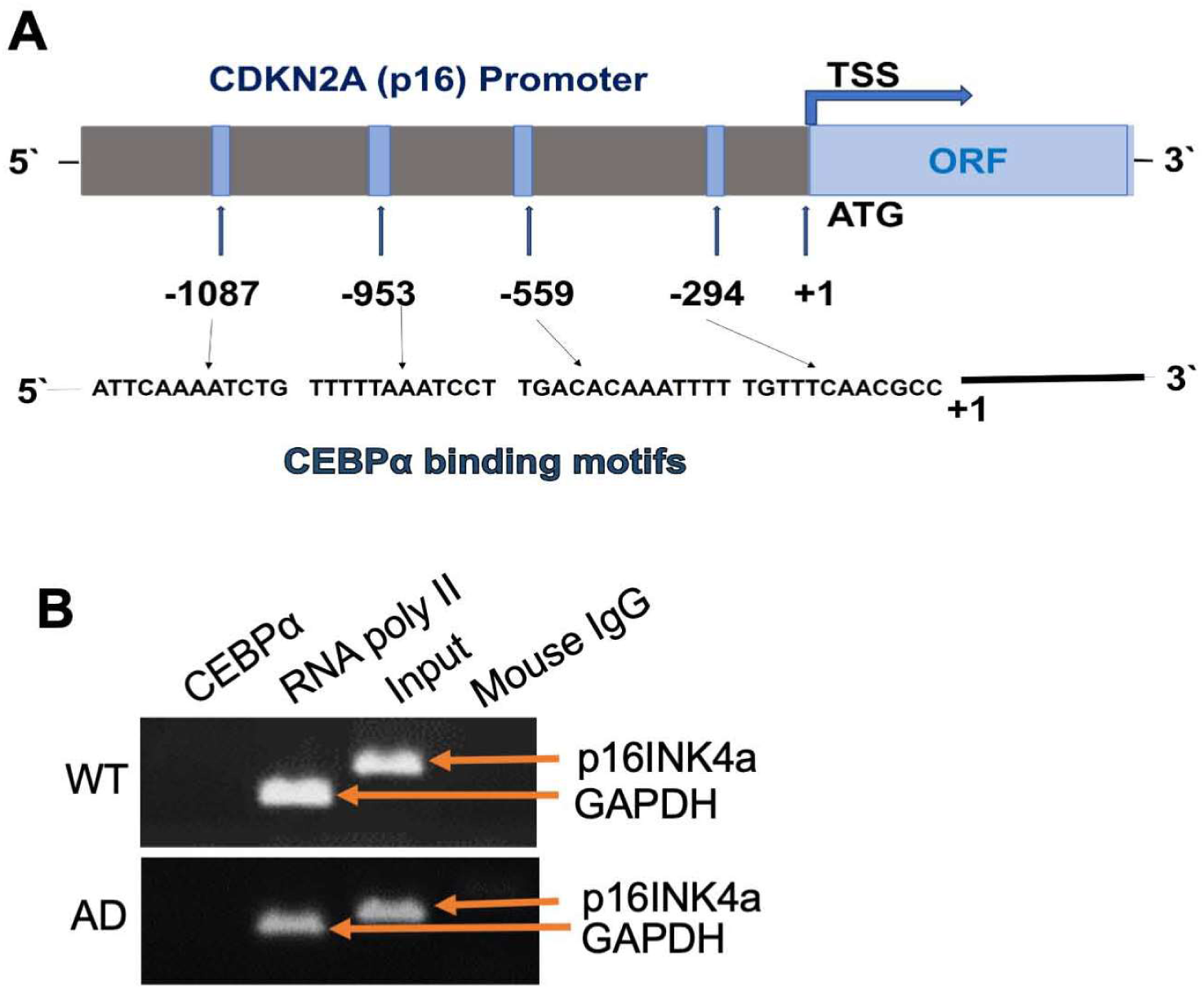
The binding of CEBPα to the promoter region of pl6INK4a was not detected. **(A)** Schematic diagram of the *pl6Ink4a* promoter with four CEBPα-binding sites. **(B)** Chromatin immunoprecipitation (ChIP) assay on the promoter of *pl6Ink4a* in isolated BMAds from *APP-PS1* mice (AD) and wild-type littermates (WT). CEBPα antibody or IgG antibody (negative control) were used for the ChIP assay. ChIP and input DNA were measured using RT-qPCR. RNA poly II was included as a positive control. Two independent experiments were done with the same results. Results from one experiment are shown.

**Figure S4.**
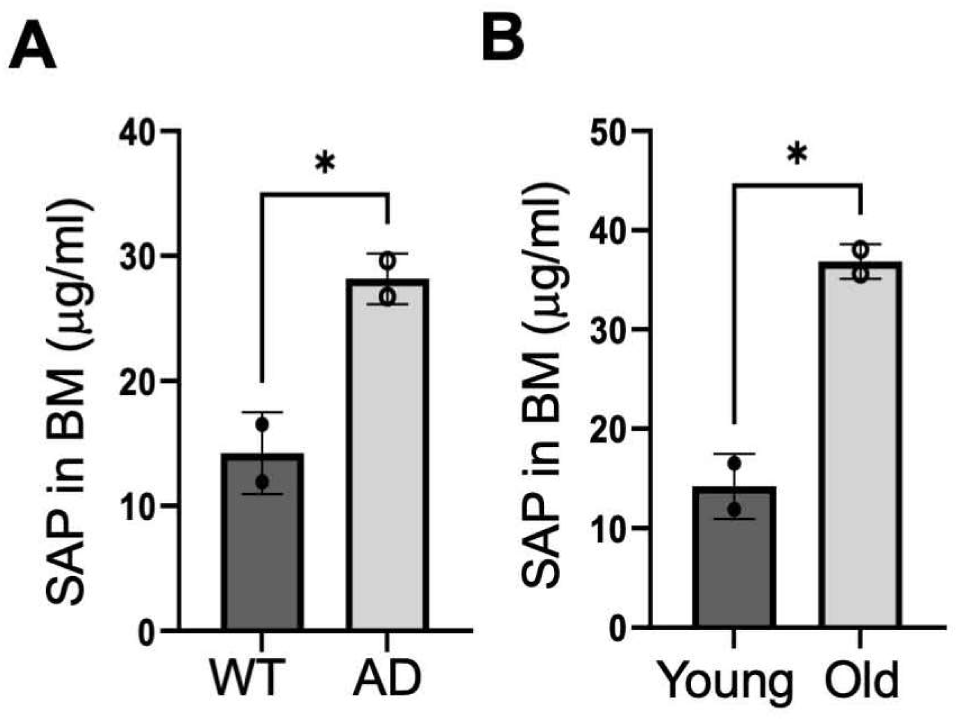
SAP concentrations are elevated in bone marrow of AD mice and aged mice. Bone marrow from long bones of AD and WT mice or from long bones of old and young mice were harvested, and the concentrations of SAP were measured. SAP concentrations are shown in (**A**) and **(B)**, respectively. Data are represented as mean ± SEM. *p < 0.05, as determined by unpaired two­tailed Student’s *t* tests for 2 groups.

**Figure S5.**
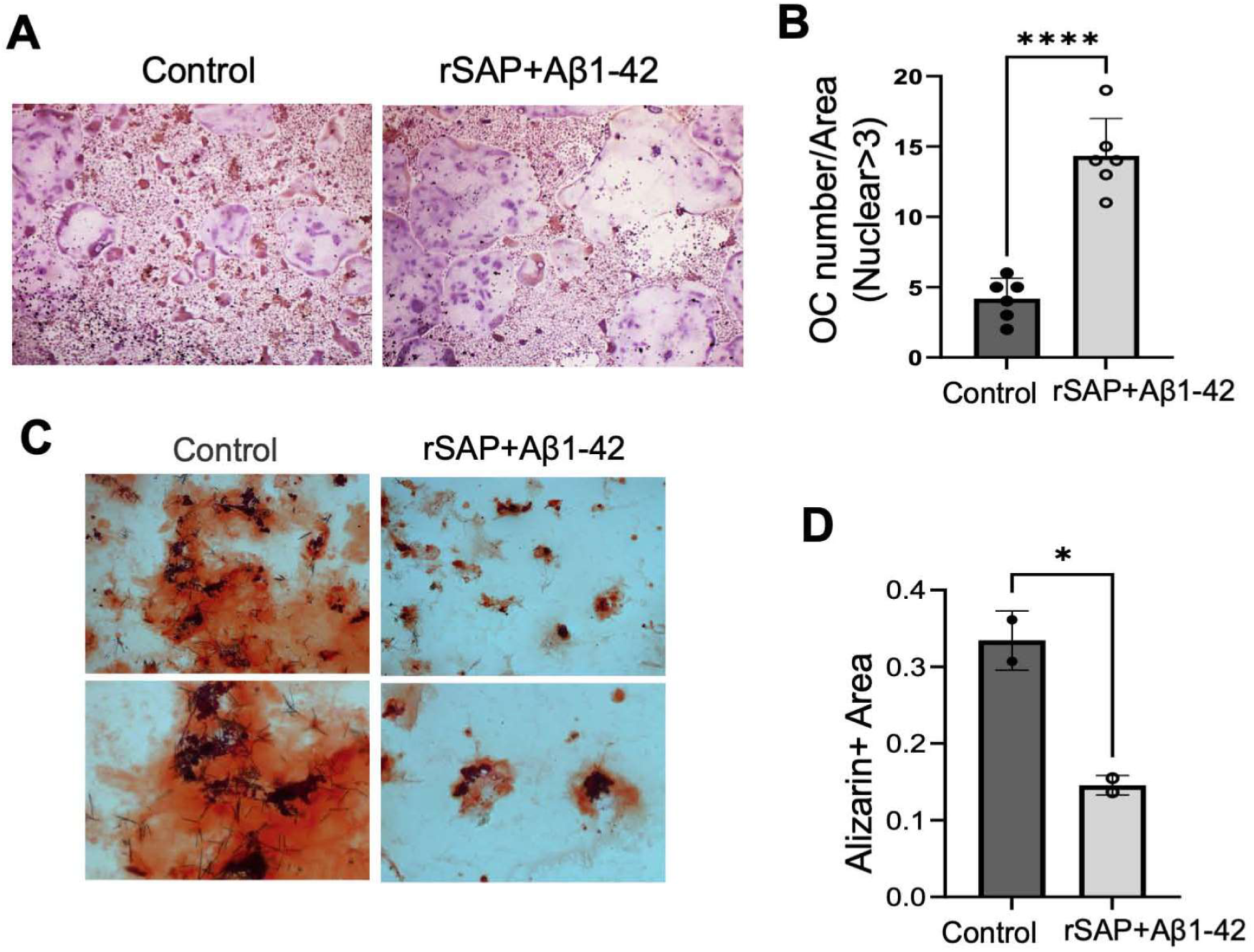
Aβ aggregates promote OCgenesis and inhibit OBgenesis. **(A-B)** Bone marrow monocytes/macrophages were cultured with 25 ng/mL M-CSF and 100 ng/mL RANKL for 3 days with or without recombinant SAP (rSAP) and Aβ1-42. Representative images of TRAP staining are shown in **(A).** Quantification of osteoclasts (>3 nucleus) per well is shown in **(B).** n= 6. **(C-D)** Mouse bone marrow mesenchymal stromal cells were cultured in osteogenic induction media with or without rSAP and Aβ1-42 for 21 days. Alizarin red staining was performed to measure the mineralization nodule formation. Representative images are shown in (C). Quantitative analysis of Alizarin red^+^ area is shown in **(D).** Data are represented as mean ± SEM. *p < 0.05, ****p < 0.0001, as determined by unpaired two-tailed Student’s *t* tests for 2 groups.

**Figure S6.**
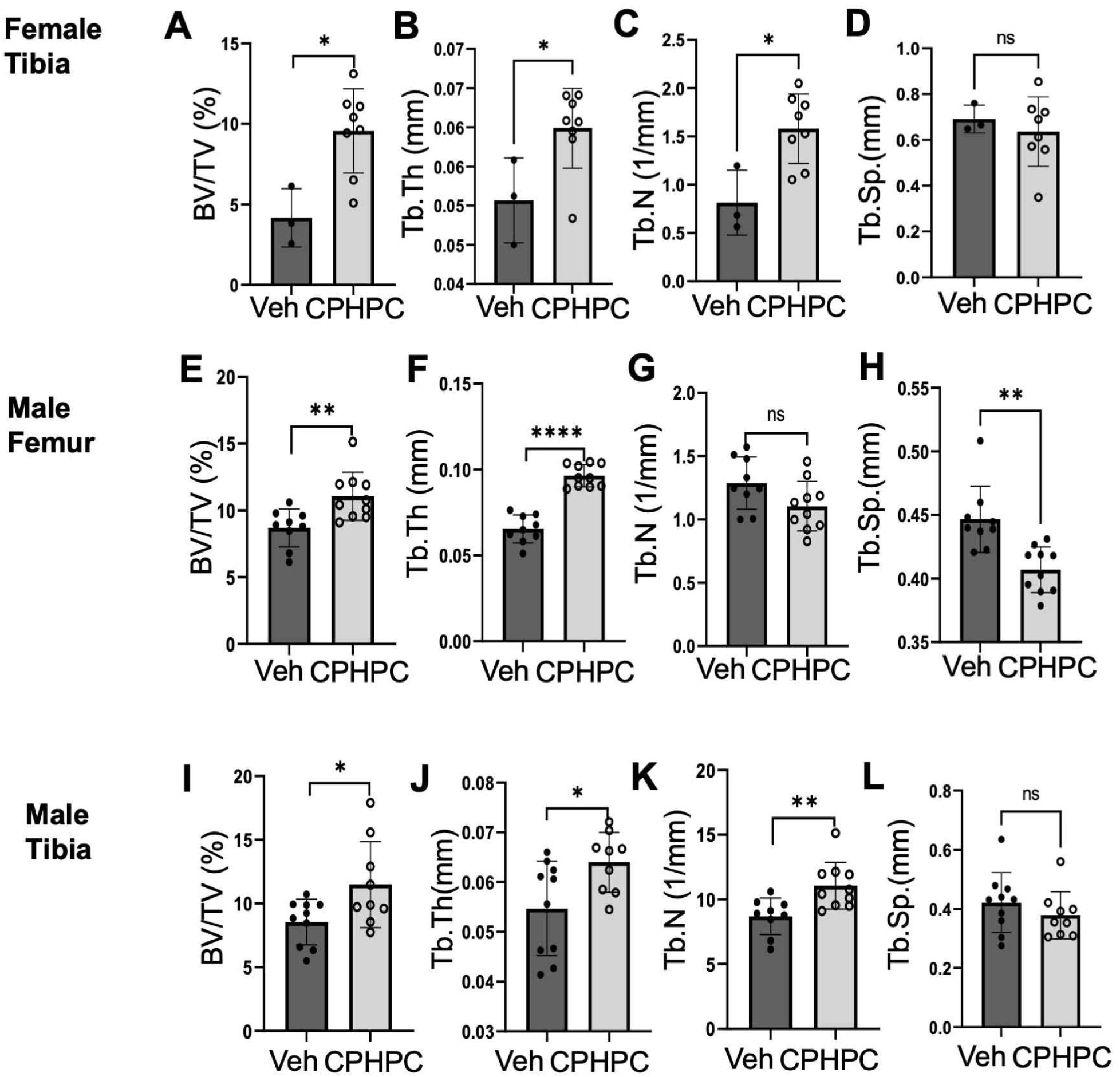
Depleting SAP by CPHPC (miridesap) increases bone mass in aged mice. Eighteen-month-old male and female C57BL/6 mice were treated with CPHPC at 5mg/ml through drinking water for 42 days. Regular drinking water was used as a control. **(A-D)** μCT analyses of the proximal tibia of female mice were performed. Quantifications of trabecular bone volume (BV/TV) **(A)**, trabecular thickness (Tb. Th) **(B)**, trabecular number (Tb. N) **(C)**, and trabecular separation (Tb. Sp) **(D)** are shown. n=5 mice. **(I-L)** μCT analyses of the proximal tibia of male mice were performed. Quantifications of trabecular bone volume (BV/TV) **(A)**, trabecular thickness (Tb. Th) **(J)**, trabecular number (Tb. N) **(K)**, and trabecular separation (Tb. Sp) (L) are shown. n=5 mice. Data are represented as mean ± SEM. *p < 0.05, **p < 0.01, ****p < 0.0001, as determined by unpaired two-tailed Student’s *t* tests for 2 groups.

## Video

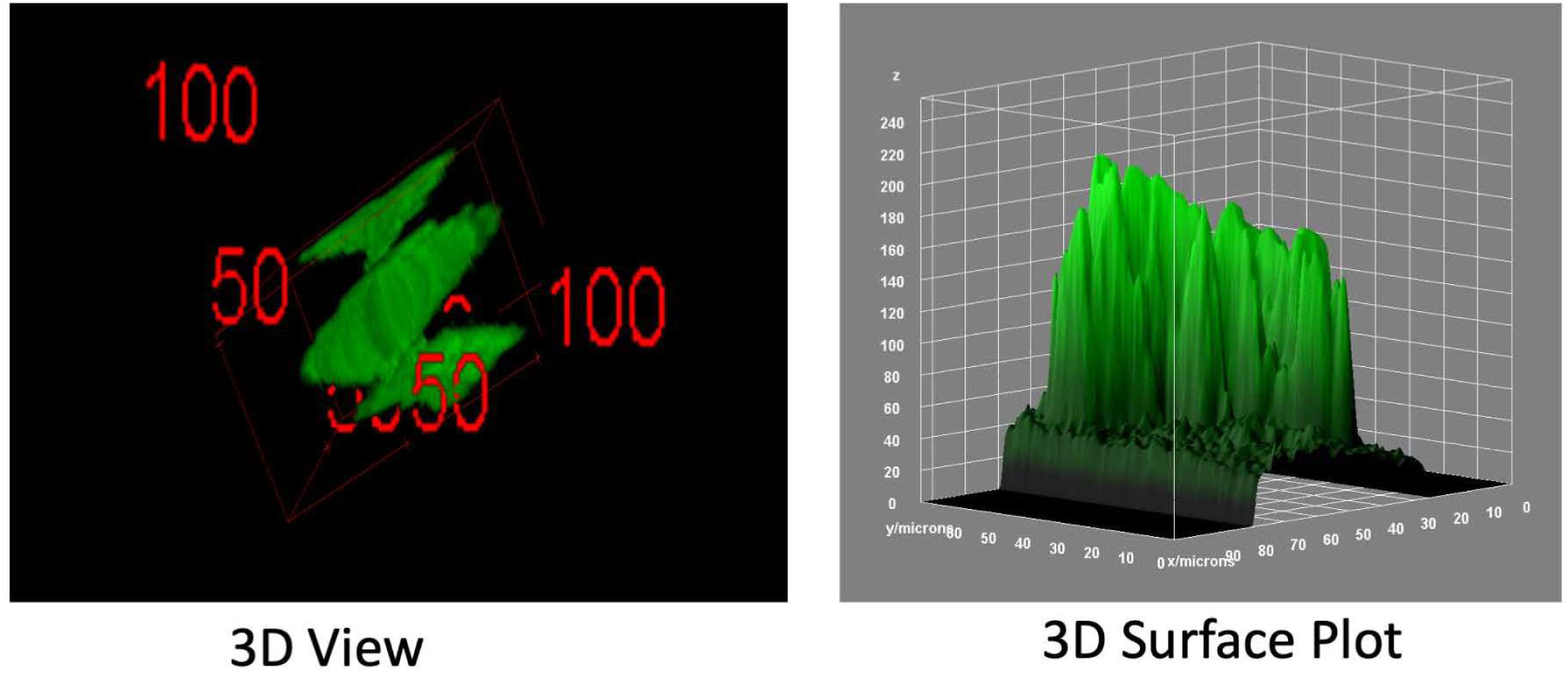

## Notes

### Competing Interest Statement

The authors have declared no competing interest.

## REFERENCES

1. Hampel, H., Hardy, J., Blennow, K., Chen, C., Perry, G., Kim, S.H., Villemagne, V.L., Aisen, P., Vendruscolo, M., Iwatsubo, T., et al. (2021). The Amyloid-beta Pathway in Alzheimer’s Disease. Mol Psychiatry 26, 5481–5503. 10.1038/s41380-021-01249-0.

2. Murphy, M.P., and LeVine, H., 3rd (2010). Alzheimer’s disease and the amyloid-beta peptide. J Alzheimers Dis 19, 311–323. 10.3233/JAD-2010-1221.

3. Cras, P., Kawai, M., Lowery, D., Gonzalez-DeWhitt, P., Greenberg, B., and Perry, G. (1991). Senile plaque neurites in Alzheimer disease accumulate amyloid precursor protein. Proc Natl Acad Sci U S A 88, 7552–7556. 10.1073/pnas.88.17.7552.

4. Benson, M.D., Buxbaum, J.N., Eisenberg, D.S., Merlini, G., Saraiva, M.J.M., Sekijima, Y., Sipe, J.D., and Westermark, P. (2020). Amyloid nomenclature 2020: update and recommendations by the International Society of Amyloidosis (ISA) nomenclature committee. Amyloid 27, 217–222. 10.1080/13506129.2020.1835263.

5. Gertz, M.A., Comenzo, R., Falk, R.H., Fermand, J.P., Hazenberg, B.P., Hawkins, P.N., Merlini, G., Moreau, P., Ronco, P., Sanchorawala, V., et al. (2005). Definition of organ involvement and treatment response in immunoglobulin light chain amyloidosis (AL): a consensus opinion from the 10th International Symposium on Amyloid and Amyloidosis, Tours, France, 18-22 April 2004. Am J Hematol 79, 319–328. 10.1002/ajh.20381.

6. Koike, H., Misu, K., Sugiura, M., Iijima, M., Mori, K., Yamamoto, M., Hattori, N., Mukai, E., Ando, Y., Ikeda, S., and Sobue, G. (2004). Pathology of early- vs late-onset TTR Met30 familial amyloid polyneuropathy. Neurology 63, 129–138. 10.1212/01.wnl.0000132966.36437.12.

7. Gertz, M.A., and Dispenzieri, A. (2020). Systemic Amyloidosis Recognition, Prognosis, and Therapy: A Systematic Review. JAMA 324, 79–89. 10.1001/jama.2020.5493.

8. Hipp, M.S., Kasturi, P., and Hartl, F.U. (2019). The proteostasis network and its decline in ageing. Nat Rev Mol Cell Biol 20, 421–435. 10.1038/s41580-019-0101-y.

9. Chiti, F., and Dobson, C.M. (2006). Protein misfolding, functional amyloid, and human disease. Annu Rev Biochem 75, 333–366. 10.1146/annurev.biochem.75.101304.123901.

10. Fandrich, M. (2007). On the structural definition of amyloid fibrils and other polypeptide aggregates. Cell Mol Life Sci 64, 2066–2078. 10.1007/s00018-007-7110-2.

11. Sawaya, M.R., Sambashivan, S., Nelson, R., Ivanova, M.I., Sievers, S.A., Apostol, M.I., Thompson, M.J., Balbirnie, M., Wiltzius, J.J., McFarlane, H.T., et al. (2007). Atomic structures of amyloid cross-beta spines reveal varied steric zippers. Nature 447, 453–457. 10.1038/nature05695.

12. Buxbaum, J.N., Dispenzieri, A., Eisenberg, D.S., Fandrich, M., Merlini, G., Saraiva, M.J.M., Sekijima, Y., and Westermark, P. (2022). Amyloid nomenclature 2022: update, novel proteins, and recommendations by the International Society of Amyloidosis (ISA) Nomenclature Committee. Amyloid 29, 213–219. 10.1080/13506129.2022.2147636.

13. Goldis, R., Kaplan, B., Kukuy, O.L., Arad, M., Magen, H., Shavit-Stein, E., Dori, A., and Livneh, A. (2023). Diagnostic Challenges and Solutions in Systemic Amyloidosis. Int J Mol Sci 24. 10.3390/ijms24054655.

14. Li, Y., Dai, J., Kametani, F., Yazaki, M., Ishigami, A., Mori, M., Miyahara, H., and Higuchi, K. (2023). Renal Function in Aged C57BL/6J Mice Is Impaired by Deposition of Age-Related Apolipoprotein A-II Amyloid Independent of Kidney Aging. Am J Pathol 193, 725–739. 10.1016/j.ajpath.2023.03.002.

15. Simons, M., Keller, P., De Strooper, B., Beyreuther, K., Dotti, C.G., and Simons, K. (1998). Cholesterol depletion inhibits the generation of beta-amyloid in hippocampal neurons. Proc Natl Acad Sci U S A 95, 6460–6464. 10.1073/pnas.95.11.6460.

16. Sachse, C., Fandrich, M., and Grigorieff, N. (2008). Paired beta-sheet structure of an Abeta(1-40) amyloid fibril revealed by electron microscopy. Proc Natl Acad Sci U S A 105, 7462–7466. 10.1073/pnas.0712290105.

17. Luhrs, T., Ritter, C., Adrian, M., Riek-Loher, D., Bohrmann, B., Dobeli, H., Schubert, D., and Riek, R. (2005). 3D structure of Alzheimer’s amyloid-beta(1-42) fibrils. Proc Natl Acad Sci U S A 102, 17342–17347. 10.1073/pnas.0506723102.

18. Glabe, C.G. (2008). Structural classification of toxic amyloid oligomers. J Biol Chem 283, 29639–29643. 10.1074/jbc.R800016200.

19. Haass, C., and Selkoe, D.J. (2007). Soluble protein oligomers in neurodegeneration: lessons from the Alzheimer’s amyloid beta-peptide. Nat Rev Mol Cell Biol 8, 101–112. 10.1038/nrm2101.

20. Cathcart, E.S., Comerford, F.R., and Cohen, A.S. (1965). Immunologic Studies on a Protein Extracted from Human Secondary Amyloid. N Engl J Med 273, 143–146. 10.1056/NEJM196507152730306.

21. Cathcart, E.S., Wollheim, F.A., and Cohen, A.S. (1967). Plasma protein constituents of amyloid fibrils. J Immunol 99, 376–385.

22. Connor, T., and Hole, S. (2024). Musculoskeletal Amyloidosis Imaging. In StatPearls.

23. Feitosa, V.A., Neves, P., Jorge, L.B., Noronha, I.L., and Onuchic, L.F. (2022). Renal amyloidosis: a new time for a complete diagnosis. Braz J Med Biol Res 55, e12284. 10.1590/1414-431X2022e12284.

24. Pilling, D., and Gomer, R.H. (2018). The Development of Serum Amyloid P as a Possible Therapeutic. Front Immunol 9, 2328. 10.3389/fimmu.2018.02328.

25. Frame, G., Bretland, K.A., and Dengler-Crish, C.M. (2020). Mechanistic complexities of bone loss in Alzheimer’s disease: a review. Connect Tissue Res 61, 4–18. 10.1080/03008207.2019.1624734.

26. Tan, Z.S., Seshadri, S., Beiser, A., Zhang, Y., Felson, D., Hannan, M.T., Au, R., Wolf, P.A., and Kiel, D.P. (2005). Bone mineral density and the risk of Alzheimer disease. Arch Neurol 62, 107–111. 10.1001/archneur.62.1.107.

27. Zhou, R., Zhou, H., Rui, L., and Xu, J. (2014). Bone loss and osteoporosis are associated with conversion from mild cognitive impairment to Alzheimer’s disease. Curr Alzheimer Res 11, 706–713. 10.2174/1567205011666140812115818.

28. Amouzougan, A., Lafaie, L., Marotte, H., Denarie, D., Collet, P., Pallot-Prades, B., and Thomas, T. (2017). High prevalence of dementia in women with osteoporosis. Joint Bone Spine 84, 611–614. 10.1016/j.jbspin.2016.08.002.

29. Zhou, R., Deng, J., Zhang, M., Zhou, H.D., and Wang, Y.J. (2011). Association between bone mineral density and the risk of Alzheimer’s disease. J Alzheimers Dis 24, 101–108. 10.3233/JAD-2010-101467.

30. Dengler-Crish, C.M., Ball, H.C., Lin, L., Novak, K.M., and Cooper, L.N. (2018). Evidence of Wnt/beta-catenin alterations in brain and bone of a tauopathy mouse model of Alzheimer’s disease. Neurobiol Aging 67, 148–158. 10.1016/j.neurobiolaging.2018.03.021.

31. Dengler-Crish, C.M., Smith, M.A., and Wilson, G.N. (2017). Early Evidence of Low Bone Density and Decreased Serotonergic Synthesis in the Dorsal Raphe of a Tauopathy Model of Alzheimer’s Disease. J Alzheimers Dis 55, 1605–1619. 10.3233/JAD-160658.

32. Mihara, S., Kawai, S., Gondo, T., and Ishihara, T. (1994). Intervertebral disc amyloidosis: histochemical, immunohistochemical and ultrastructural observations. Histopathology 25, 415–420. 10.1111/j.1365-2559.1994.tb00002.x.

33. Yanagisawa, A., Ueda, M., Sueyoshi, T., Nakamura, E., Tasaki, M., Suenaga, G., Motokawa, H., Toyoshima, R., Kinoshita, Y., Misumi, Y., et al. (2016). Knee osteoarthritis associated with different kinds of amyloid deposits and the impact of aging on type of amyloid. Amyloid 23, 26–32. 10.3109/13506129.2015.1115758.

34. Yanagisawa, A., Ueda, M., Sueyoshi, T., Okada, T., Fujimoto, T., Ogi, Y., Kitagawa, K., Tasaki, M., Misumi, Y., Oshima, T., et al. (2015). Amyloid deposits derived from transthyretin in the ligamentum flavum as related to lumbar spinal canal stenosis. Mod Pathol 28, 201–207. 10.1038/modpathol.2014.102.

35. Agostini, A., Yuchun, D., Li, B., Kendall, D.A., and Pardon, M.C. (2020). Sex-specific hippocampal metabolic signatures at the onset of systemic inflammation with lipopolysaccharide in the APPswe/PS1dE9 mouse model of Alzheimer’s disease. Brain Behav Immun 83, 87–111. 10.1016/j.bbi.2019.09.019.

36. Saito, T., Hisahara, S., Iwahara, N., Emoto, M.C., Yokokawa, K., Suzuki, H., Manabe, T., Matsumura, A., Suzuki, S., Matsushita, T., et al. (2019). Early administration of galantamine from preplaque phase suppresses oxidative stress and improves cognitive behavior in APPswe/PS1dE9 mouse model of Alzheimer’s disease. Free Radic Biol Med 145, 20–32. 10.1016/j.freeradbiomed.2019.09.014.

37. Hong, J.H., Kang, J.W., Kim, D.K., Baik, S.H., Kim, K.H., Shanta, S.R., Jung, J.H., Mook-Jung, I., and Kim, K.P. (2016). Global changes of phospholipids identified by MALDI imaging mass spectrometry in a mouse model of Alzheimer’s disease. J Lipid Res 57, 36–45. 10.1194/jlr.M057869.

38. Oakley, H., Cole, S.L., Logan, S., Maus, E., Shao, P., Craft, J., Guillozet-Bongaarts, A., Ohno, M., Disterhoft, J., Van Eldik, L., et al. (2006). Intraneuronal beta-amyloid aggregates, neurodegeneration, and neuron loss in transgenic mice with five familial Alzheimer’s disease mutations: potential factors in amyloid plaque formation. J Neurosci 26, 10129–10140. 10.1523/JNEUROSCI.1202-06.2006.

39. Ali, D., Tencerova, M., Figeac, F., Kassem, M., and Jafari, A. (2022). The pathophysiology of osteoporosis in obesity and type 2 diabetes in aging women and men: The mechanisms and roles of increased bone marrow adiposity. Front Endocrinol (Lausanne) 13, 981487. 10.3389/fendo.2022.981487.

40. Armas, L.A., and Recker, R.R. (2012). Pathophysiology of osteoporosis: new mechanistic insights. Endocrinol Metab Clin North Am 41, 475–486. 10.1016/j.ecl.2012.04.006.

41. Paranthaman, M., Angu Bala Ganesh, K.S.V., and Silambanan, S. (2024). Linking bone marrow fat with decreased bone mineral density among Indian patients with osteoporotic fracture. Bioinformation 20, 49–54. 10.6026/973206300200049.

42. Liu, X., Gu, Y., Kumar, S., Amin, S., Guo, Q., Wang, J., Fang, C.L., Cao, X., and Wan, M. (2023). Oxylipin-PPARgamma-initiated adipocyte senescence propagates secondary senescence in the bone marrow. Cell Metab 35, 667–684 e666. 10.1016/j.cmet.2023.03.005.

43. Kang, M.A., So, E.Y., Simons, A.L., Spitz, D.R., and Ouchi, T. (2012). DNA damage induces reactive oxygen species generation through the *H2AX*-Nox1/Rac1 pathway. Cell Death Dis 3, e249. 10.1038/cddis.2011.134.

44. Katsuta, E., Sawant Dessai, A., Ebos, J.M., Yan, L., Ouchi, T., and Takabe, K. (2022). *H2AX* mRNA expression reflects DNA repair, cell proliferation, metastasis, and worse survival in breast cancer. Am J Cancer Res 12, 793–804.

45. Aquino-Martinez, R., Rowsey, J.L., Fraser, D.G., Eckhardt, B.A., Khosla, S., Farr, J.N., and Monroe, D.G. (2020). LPS-induced premature osteocyte senescence: Implications in inflammatory alveolar bone loss and periodontal disease pathogenesis. Bone 132, 115220. 10.1016/j.bone.2019.115220.

46. Farr, J.N., Xu, M., Weivoda, M.M., Monroe, D.G., Fraser, D.G., Onken, J.L., Negley, B.A., Sfeir, J.G., Ogrodnik, M.B., Hachfeld, C.M., et al. (2017). Targeting cellular senescence prevents age-related bone loss in mice. Nat Med 23, 1072–1079. 10.1038/nm.4385.

47. Farr, J.N., Fraser, D.G., Wang, H., Jaehn, K., Ogrodnik, M.B., Weivoda, M.M., Drake, M.T., Tchkonia, T., LeBrasseur, N.K., Kirkland, J.L., et al. (2016). Identification of Senescent Cells in the Bone Microenvironment. J Bone Miner Res 31, 1920–1929. 10.1002/jbmr.2892.

48. Gan, Q., Huang, J., Zhou, R., Niu, J., Zhu, X., Wang, J., Zhang, Z., and Tong, T. (2008). PPARgamma accelerates cellular senescence by inducing *p16Ink4a*lpha expression in human diploid fibroblasts. J Cell Sci 121, 2235–2245. 10.1242/jcs.026633.

49. Takasugi, M., Yoshida, Y., and Ohtani, N. (2022). Cellular senescence and the tumour microenvironment. Mol Oncol 16, 3333–3351. 10.1002/1878-0261.13268.

50. Coppe, J.P., Patil, C.K., Rodier, F., Sun, Y., Munoz, D.P., Goldstein, J., Nelson, P.S., Desprez, P.Y., and Campisi, J. (2008). Senescence-associated secretory phenotypes reveal cell-nonautonomous functions of oncogenic RAS and the p53 tumor suppressor. PLoS Biol 6, 2853–2868. 10.1371/journal.pbio.0060301.

51. Kuilman, T., Michaloglou, C., Vredeveld, L.C., Douma, S., van Doorn, R., Desmet, C.J., Aarden, L.A., Mooi, W.J., and Peeper, D.S. (2008). Oncogene-induced senescence relayed by an interleukin-dependent inflammatory network. Cell 133, 1019–1031. 10.1016/j.cell.2008.03.039.

52. Acosta, J.C., O’Loghlen, A., Banito, A., Guijarro, M.V., Augert, A., Raguz, S., Fumagalli, M., Da Costa, M., Brown, C., Popov, N., et al. (2008). Chemokine signaling via the CXCR2 receptor reinforces senescence. Cell 133, 1006–1018. 10.1016/j.cell.2008.03.038.

53. Wu, W., Fu, J., Gu, Y., Wei, Y., Ma, P., and Wu, J. (2020). JAK2/STAT3 regulates estrogen-related senescence of bone marrow stem cells. J Endocrinol 245, 141–153. 10.1530/JOE-19-0518.

54. van Deursen, J.M. (2014). The role of senescent cells in ageing. Nature 509, 439–446. 10.1038/nature13193.

55. Klotz, S.A., and Lipke, P.N. (2022). The Paradoxical Effects of Serum Amyloid-β Component on Disseminated Candidiasis. Pathogens 11. 10.3390/pathogens11111304.

56. Pepys, M.B. (2018). The Pentraxins 1975-2018: Serendipity, Diagnostics and Drugs. Front Immunol 9, 2382. 10.3389/fimmu.2018.02382.

57. Pepys, M.B., Herbert, J., Hutchinson, W.L., Tennent, G.A., Lachmann, H.J., Gallimore, J.R., Lovat, L.B., Bartfai, T., Alanine, A., Hertel, C., et al. (2002). Targeted pharmacological depletion of serum amyloid P component for treatment of human amyloidosis. Nature 417, 254–259. 10.1038/417254a.

58. Klotz, S.A., Sobonya, R.E., Lipke, P.N., and Garcia-Sherman, M.C. (2016). Serum Amyloid P Component and Systemic Fungal Infection: Does It Protect the Host or Is It a Trojan Horse? Open Forum Infect Dis 3, ofw166. 10.1093/ofid/ofw166.

59. Gilchrist, K.B., Garcia, M.C., Sobonya, R., Lipke, P.N., and Klotz, S.A. (2012). New features of invasive candidiasis in humans: amyloid formation by fungi and deposition of serum amyloid P component by the host. J Infect Dis 206, 1473–1478. 10.1093/infdis/jis464.

60. Pepys, M.B. (1999). Serum amyloid P component (not serum amyloid protein). Nat Med 5, 852–853. 10.1038/11272.

61. Pepys, M.B., Dyck, R.F., de Beer, F.C., Skinner, M., and Cohen, A.S. (1979). Binding of serum amyloid P-component (SAP) by amyloid fibrils. Clin Exp Immunol 38, 284–293.

62. Victorelli, S., Salmonowicz, H., Chapman, J., Martini, H., Vizioli, M.G., Riley, J.S., Cloix, C., Hall-Younger, E., Machado Espindola-Netto, J., Jurk, D., et al. (2023). Apoptotic stress causes mtDNA release during senescence and drives the SASP. Nature 622, 627–636. 10.1038/s41586-023-06621-4.

63. Saul, D., Kosinsky, R.L., Atkinson, E.J., Doolittle, M.L., Zhang, X., LeBrasseur, N.K., Pignolo, R.J., Robbins, P.D., Niedernhofer, L.J., Ikeno, Y., et al. (2022). A new gene set identifies senescent cells and predicts senescence-associated pathways across tissues. Nat Commun 13, 4827. 10.1038/s41467-022-32552-1.

64. Kudlova, N., De Sanctis, J.B., and Hajduch, M. (2022). Cellular Senescence: Molecular Targets, Biomarkers, and Senolytic Drugs. Int J Mol Sci 23. 10.3390/ijms23084168.

65. Mohamad Kamal, N.S., Safuan, S., Shamsuddin, S., and Foroozandeh, P. (2020). Aging of the cells: Insight into cellular senescence and detection Methods. Eur J Cell Biol 99, 151108. 10.1016/j.ejcb.2020.151108.

66. Farr, J.N., and Khosla, S. (2019). Cellular senescence in bone. Bone 121, 121–133. 10.1016/j.bone.2019.01.015.

67. Xu, M., Pirtskhalava, T., Farr, J.N., Weigand, B.M., Palmer, A.K., Weivoda, M.M., Inman, C.L., Ogrodnik, M.B., Hachfeld, C.M., Fraser, D.G., et al. (2018). Senolytics improve physical function and increase lifespan in old age. Nat Med 24, 1246–1256. 10.1038/s41591-018-0092-9.

68. Tiwari, S., Atluri, V., Kaushik, A., Yndart, A., and Nair, M. (2019). Alzheimer’s disease: pathogenesis, diagnostics, and therapeutics. Int J Nanomedicine 14, 5541–5554. 10.2147/IJN.S200490.

69. Weller, J., and Budson, A. (2018). Current understanding of Alzheimer’s disease diagnosis and treatment. F1000Res 7. 10.12688/f1000research.14506.1.

70. Xia, W.F., Jung, J.U., Shun, C., Xiong, S., Xiong, L., Shi, X.M., Mei, L., and Xiong, W.C. (2013). Swedish mutant APP suppresses osteoblast differentiation and causes osteoporotic deficit, which are ameliorated by N-acetyl-L-cysteine. J Bone Miner Res 28, 2122–2135. 10.1002/jbmr.1954.

71. Lee, L.Y., Chou, W., Chen, W.P., Wang, M.F., Chen, Y.J., Chen, C.C., and Tung, K.C. (2021). Erinacine A-Enriched Hericium erinaceus Mycelium Delays Progression of Age-Related Cognitive Decline in Senescence Accelerated Mouse Prone 8 (SAMP8) Mice. Nutrients 13. 10.3390/nu13103659.

72. Malerba, H.N., Pereira, A.A.R., Pierrobon, M.F., Abrao, G.S., Toricelli, M., Akamine, E.H., Buck, H.S., and Viel, T.A. (2021). Combined Neuroprotective Strategies Blocked Neurodegeneration and Improved Brain Function in Senescence-Accelerated Mice. Front Aging Neurosci 13, 681498. 10.3389/fnagi.2021.681498.

73. Cheng, X.R., Zhou, W.X., and Zhang, Y.X. (2014). The behavioral, pathological and therapeutic features of the senescence-accelerated mouse prone 8 strain as an Alzheimer’s disease animal model. Ageing Res Rev 13, 13–37. 10.1016/j.arr.2013.10.002.

74. Piccarducci, R., Pietrobono, D., Pellegrini, C., Daniele, S., Fornai, M., Antonioli, L., Trincavelli, M.L., Blandizzi, C., and Martini, C. (2019). High Levels of beta-Amyloid, Tau, and Phospho-Tau in Red Blood Cells as Biomarkers of Neuropathology in Senescence-Accelerated Mouse. Oxid Med Cell Longev 2019, 5030475. 10.1155/2019/5030475.

75. Butterfield, D.A., and Poon, H.F. (2005). The senescence-accelerated prone mouse (SAMP8): a model of age-related cognitive decline with relevance to alterations of the gene expression and protein abnormalities in Alzheimer’s disease. Exp Gerontol 40, 774–783. 10.1016/j.exger.2005.05.007.

76. Pilling, D., Cox, N., Thomson, M.A., Karhadkar, T.R., and Gomer, R.H. (2019). Serum Amyloid P and a Dendritic Cell-Specific Intercellular Adhesion Molecule-3-Grabbing Nonintegrin Ligand Inhibit High-Fat Diet-Induced Adipose Tissue and Liver Inflammation and Steatosis in Mice. Am J Pathol 189, 2400–2413. 10.1016/j.ajpath.2019.08.005.

77. Hutchinson, W.L., Hohenester, E., and Pepys, M.B. (2000). Human serum amyloid P component is a single uncomplexed pentamer in whole serum. Mol Med 6, 482–493.

78. Iwanaga, T., Wakasugi, S., Inomoto, T., Uehira, M., Ohnishi, S., Nishiguchi, S., Araki, K., Uno, M., Miyazaki, J., Maeda, S., and, et al. (1989). Liver-specific and high-level expression of human serum amyloid P component gene in transgenic mice. Dev Genet 10, 365–371. 10.1002/dvg.1020100504.

79. Pepys, M.B., Booth, D.R., Hutchinson, W.L., Gallimore, J.R., Collins, I.M., and Hohenester, E. (1997). Amyloid P component. A critical review. Amyloid 4, 274–295. 10.3109/13506129709003838.

80. Tennent, G.A., Lovat, L.B., and Pepys, M.B. (1995). Serum amyloid P component prevents proteolysis of the amyloid fibrils of Alzheimer disease and systemic amyloidosis. Proc Natl Acad Sci U S A 92, 4299–4303. 10.1073/pnas.92.10.4299.

81. Mold, M., Shrive, A.K., and Exley, C. (2012). Serum amyloid P component accelerates the formation and enhances the stability of amyloid fibrils in a physiologically significant under-saturated solution of amyloid-beta42. J Alzheimers Dis 29, 875–881. 10.3233/JAD-2012-120076.

82. Hamazaki, H. (1995). Amyloid P component promotes aggregation of Alzheimer’s beta-amyloid peptide. Biochem Biophys Res Commun 211, 349–353. 10.1006/bbrc.1995.1819.

83. Botto, M., Hawkins, P.N., Bickerstaff, M.C., Herbert, J., Bygrave, A.E., McBride, A., Hutchinson, W.L., Tennent, G.A., Walport, M.J., and Pepys, M.B. (1997). Amyloid deposition is delayed in mice with targeted deletion of the serum amyloid P component gene. Nat Med 3, 855–859. 10.1038/nm0897-855.

84. Al-Shawi, R., Tennent, G.A., Millar, D.J., Richard-Londt, A., Brandner, S., Werring, D.J., Simons, J.P., and Pepys, M.B. (2016). Pharmacological removal of serum amyloid P component from intracerebral plaques and cerebrovascular Abeta amyloid deposits in vivo. Open Biol 6, 150202. 10.1098/rsob.150202.

85. Schmidt, A.F., Finan, C., Chopade, S., Ellmerich, S., Rossor, M.N., Hingorani, A.D., and Pepys, M.B. (2023). Genetic evidence for serum amyloid P component as a drug target for treatment of neurodegenerative disorders. medRxiv. 10.1101/2023.08.15.23293564.

86. Richards, D.B., Cookson, L.M., Barton, S.V., Liefaard, L., Lane, T., Hutt, D.F., Ritter, J.M., Fontana, M., Moon, J.C., Gillmore, J.D., et al. (2018). Repeat doses of antibody to serum amyloid P component clear amyloid deposits in patients with systemic amyloidosis. Sci Transl Med 10. 10.1126/scitranslmed.aan3128.

87. Sahota, T., Berges, A., Barton, S., Cookson, L., Zamuner, S., and Richards, D. (2015). Target Mediated Drug Disposition Model of CPHPC in Patients with Systemic Amyloidosis. CPT Pharmacometrics Syst Pharmacol 4, e15. 10.1002/psp4.15.

88. Li, T., Braunstein, K.E., Zhang, J., Lau, A., Sibener, L., Deeble, C., and Wong, P.C. (2016). The neuritic plaque facilitates pathological conversion of tau in an Alzheimer’s disease mouse model. Nat Commun 7, 12082. 10.1038/ncomms12082.

89. LaClair, K.D., Manaye, K.F., Lee, D.L., Allard, J.S., Savonenko, A.V., Troncoso, J.C., and Wong, P.C. (2013). Treatment with bexarotene, a compound that increases apolipoprotein-E, provides no cognitive benefit in mutant APP/PS1 mice. Mol Neurodegener 8, 18. 10.1186/1750-1326-8-18.

90. Naruse, S., Thinakaran, G., Luo, J.J., Kusiak, J.W., Tomita, T., Iwatsubo, T., Qian, X., Ginty, D.D., Price, D.L., Borchelt, D.R., et al. (1998). Effects of PS1 deficiency on membrane protein trafficking in neurons. Neuron 21, 1213–1221. 10.1016/s0896-6273(00)80637-6.

91. Santhanam, L., Liu, G., Jandu, S., Su, W., Wodu, B.P., Savage, W., Poe, A., Liu, X., Alexander, L.M., Cao, X., and Wan, M. (2021). Skeleton-secreted PDGF-BB mediates arterial stiffening. J Clin Invest 131. 10.1172/JCI147116.

92. Li, C., Xing, Q., Yu, B., Xie, H., Wang, W., Shi, C., Crane, J.L., Cao, X., and Wan, M. (2013). Disruption of LRP6 in osteoblasts blunts the bone anabolic activity of PTH. J Bone Miner Res 28, 2094–2108. 10.1002/jbmr.1962.

93. Hu, B., Lv, X., Chen, H., Xue, P., Gao, B., Wang, X., Zhen, G., Crane, J.L., Pan, D., Liu, S., et al. (2020). Sensory nerves regulate mesenchymal stromal cell lineage commitment by tuning sympathetic tones. J Clin Invest 130, 3483–3498. 10.1172/JCI131554.

94. Fan, Y., Hanai, J.I., Le, P.T., Bi, R., Maridas, D., DeMambro, V., Figueroa, C.A., Kir, S., Zhou, X., Mannstadt, M., et al. (2017). Parathyroid Hormone Directs Bone Marrow Mesenchymal Cell Fate. Cell Metab 25, 661–672. 10.1016/j.cmet.2017.01.001.

95. Scheller, E.L., Doucette, C.R., Learman, B.S., Cawthorn, W.P., Khandaker, S., Schell, B., Wu, B., Ding, S.Y., Bredella, M.A., Fazeli, P.K., et al. (2015). Region-specific variation in the properties of skeletal adipocytes reveals regulated and constitutive marrow adipose tissues. Nat Commun 6, 7808. 10.1038/ncomms8808.

96. Su, W., Liu, G., Mohajer, B., Wang, J., Shen, A., Zhang, W., Liu, B., Guermazi, A., Gao, P., Cao, X., et al. (2022). Senescent preosteoclast secretome promotes metabolic syndrome associated osteoarthritis through cyclooxygenase 2. Elife 11. 10.7554/eLife.79773.

97. Wang, J., Fang, C.L., Noller, K., Wei, Z., Liu, G., Shen, K., Song, K., Cao, X., and Wan, M. (2023). Bone-derived PDGF-BB drives brain vascular calcification in male mice. J Clin Invest 133. 10.1172/JCI168447.

98. Liu, X., Chai, Y., Liu, G., Su, W., Guo, Q., Lv, X., Gao, P., Yu, B., Ferbeyre, G., Cao, X., and Wan, M. (2021). Osteoclasts protect bone blood vessels against senescence through the angiogenin/plexin-B2 axis. Nat Commun 12, 1832. 10.1038/s41467-021-22131-1.

99. Friedrich, R.P., Tepper, K., Ronicke, R., Soom, M., Westermann, M., Reymann, K., Kaether, C., and Fandrich, M. (2010). Mechanism of amyloid plaque formation suggests an intracellular basis of Abeta pathogenicity. Proc Natl Acad Sci U S A 107, 1942–1947. 10.1073/pnas.0904532106.

100. Maniv, I., Sarji, M., Bdarneh, A., Feldman, A., Ankawa, R., Koren, E., Magid-Gold, I., Reis, N., Soteriou, D., Salomon-Zimri, S., et al. (2023). Altered ubiquitin signaling induces Alzheimer’s disease-like hallmarks in a three-dimensional human neural cell culture model. Nat Commun 14, 5922. 10.1038/s41467-023-41545-7.

